# Mechanism of Co-Transcriptional Cap-Snatching by Influenza Polymerase

**DOI:** 10.1101/2024.08.11.607481

**Authors:** Alexander Helmut Rotsch, Delong Li, Maud Dupont, Tim Krischuns, Christiane Oberthuer, Alice Stelfox, Maria Lukarska, Isaac Fianu, Michael Lidschreiber, Nadia Naffakh, Christian Dienemann, Stephen Cusack, Patrick Cramer

**Affiliations:** Department of Molecular Biology, Max Planck Institute for Multidisciplinary Science, Goettingen, Germany; Mechanisms of Cellular Quality Control, Max-Planck-Institute of Biophysics, Frankfurt, Germany; Institut Pasteur, Université Paris Cité, CNRS UMR3569, RNA Biology and Influenza Viruses, Paris, France; European Molecular Biology Laboratory, Grenoble, France; Department of Molecular and Cell Biology, University of California, Berkeley, CA, USA

**Keywords:** Influenza virus, influenza A, Cap-snatching, Cryo-electron microscopy, Transcription

## Abstract

Influenza virus mRNA is stable and competent for nuclear export and translation because it receives a 5′ cap(1) structure in a process called cap-snatching^1^. During cap-snatching, the viral RNA-dependent RNA polymerase (FluPol) binds to host RNA polymerase II (Pol II) and the emerging transcript^2,3^. The FluPol endonuclease then cleaves a capped RNA fragment that subsequently acts as a primer for the transcription of viral genes^4,5^. Here, we present the cryo-EM structure of FluPol bound to a transcribing Pol II in complex with the elongation factor DSIF in the pre-cleavage state. The structure shows that FluPol directly interacts with both Pol II and DSIF, which position the FluPol endonuclease domain near the RNA exit channel of Pol II. These interactions are important for the endonuclease activity of FluPol and FluPol activity in cells. A second structure trapped after cap-snatching shows that cleavage rearranges the capped RNA primer within the FluPol, directing the capped RNA 3′-end towards the FluPol polymerase active site for viral transcription initiation. Altogether, our results provide the molecular mechanisms of co-transcriptional cap-snatching by FluPol.

## Introduction

Influenza is an acute respiratory disease that causes 290,000 to 650,000 human deaths each year^6,7^. Influenza is caused by an infection with influenza A or B viruses, which circulate in temperate regions as seasonal influenza^6^. However, rare zoonotic transmissions can cause pandemic influenza outbreaks with high mortality and economic losses^8,9^. There is current concern that the unexpected susceptibility of dairy cows to avian H5N1 strains may be path towards to a new pandemic^10–12^. Influenza viruses are segmented negative-sense RNA viruses infecting the respiratory tract epithelial cells in humans^9^. Upon infection, the eight viral ribonucleoproteins are released into the cytoplasm and imported into the nucleus, where transcription of viral genes into mRNA and replication of the viral genome occurs^13,14^. Each viral ribonucleoprotein contains a genome segment that is encapsidated by multiple copies of the viral nucleoprotein and one copy of the viral RNA-dependent RNA polymerase (FluPol). FluPol consists of subunits PA, PB1, and PB2 and has been structurally characterised^2,15,16^.

Viral transcripts must contain a 5′ cap structure and a 3′ poly-A tail to ensure stability, nuclear export, and efficient translation^17^. However, unlike non-segmented negative-sense RNA viruses, the influenza virus genome does not encode enzymes to synthesize a 5′ cap^18^. Instead, FluPol utilizes capped RNA primers that are cleaved from nascent host transcripts in a process called cap-snatching^1,5^. The FluPol PB2 cap-binding domain binds a nascent 5′ capped host RNA, and the PA endonuclease domain cleaves off 10-15 nt from the 5′ end. The 3′-terminal nucleotides of this RNA primer then anneal to the 3′ end of the viral genome segment and prime transcription of the viral mRNA^15,19,20^.

Capped host transcripts are synthesized by cellular RNA polymerase II (Pol II). Pol II transcription starts with assembling a pre-initiation complex consisting of Pol II and the general transcription factors at gene promoters^21^. To escape from the gene promoter, the largest Pol II subunit RPB1 C-terminal domain (CTD) heptad repeats are phosphorylated at serine 5 and 7 by the TFIIH CDK-activating kinase (CAK) ^22,23^. CTD phosphorylation and the growing nascent RNA transcript cause the initiation factors to dissociate from Pol II^23,24^. Recruitment of the elongation factor DSIF after synthesis of ∼20 nt of RNA establishes the early Pol II elongation complex (Pol II-DSIF EC). This complex is then converted to a paused elongation complex (PEC) containing the negative elongation factor NELF at a transcript length of 25-50 nt^24–26^. Synthesis of the 5′ cap occurs co-transcriptionally by the capping enzymes RNGTT, RNMT, and CMTR1^26^ in the context of the Pol II-DSIF EC or the PEC. RNGTT is a bifunctional enzyme acting as a triphosphatase and guanylyltransferase, creating a GpppN structure at the 5′end of the Pol II transcript. RNMT and CMTR1 are methyltransferases adding a methyl group to N7 of the cap-guanosine and the 2′-OH of the first regular nucleotide, respectively, producing the m7GpppmN cap(1) structure^26^, which the cap-binding domain of PB2 tightly binds during cap-snatching^27,28^.

Cap-snatching depends on host transcription as it has been shown that inhibition of Pol II genes as well as with the Pol II CTD that is phosphorylated at serine 5 residues, indicating that cap snatching occurs during early phases of Pol II transcription^2,29–31^. Cell-based protein-protein interaction assays indicate that FluPol does not only bind to the CTD but also the Pol II body^32^. Co-immunoprecipitation – mass spectrometry experiments have shown that the elongation factor DSIF co-purifies with FluPol^5,33^, and other studies suggest that FluPol depends on the cap(1) structure for cap-snatching^27^. However, how FluPol interacts with the host transcription machinery for cap-snatching at the molecular level is unknown.

Here, we show that FluPol binds to the transcribing Pol II-DSIF complex for efficient cap-snatching. Furthermore, we report two cryo-EM structures of FluPol bound to a Pol II-DSIF EC before and after endonucleolytic RNA cleavage by the FluPol. The structures show that during cap-snatching, the PA endonuclease domain of FluPol binds near the RNA exit channel of Pol II and that this interaction is stabilised by DSIF. Furthermore, using cell-based minigenome assays, we confirm that mutation of residues forming the interface between FluPol and the Pol II-DSIF EC reduce FluPol activity. In summary, we present the molecular mechanism of cap-snatching by FluPol.

## Results

### Cap-snatching requires an early Pol II elongation complex

To study the molecular basis of cap-snatching, we first investigated how the formation of a complex between FluPol and transcribing Pol II (Pol II EC) depends on the cap(1)-structure and CTD phosphorylation. We purified *S. scrofa* Pol II (96% identical to human Pol II) from the endogenous source^34^. Whereas in preliminary studies reconstituting the cap-snatching complex we used bat FluPol (H17N10)^31^, here we used recombinant, promoter bound FluPol from the influenza strain A/Zhejiang/DTID-ZJU01/2013(H7N9)^35,36^, (**ED Figure 1a**). To reduce RNA cleavage and enhance complex stability, we used the PA E119D mutant of FluPol (FluPol^E119D^), which has impaired endonuclease activity^37,38^. A Pol II EC containing a 35 nt cap(1)-RNA, 45 nt template, and non-template DNA was assembled as established previously^39^. The 35 nt RNA length was chosen considering a 12 nt RNA primer produced by cap-snatching^20,40^, an additional 3 nt bound by the PA endonuclease^38^, and 20 nt RNA bound within the Pol II EC^34^.

**Fig. 1.**
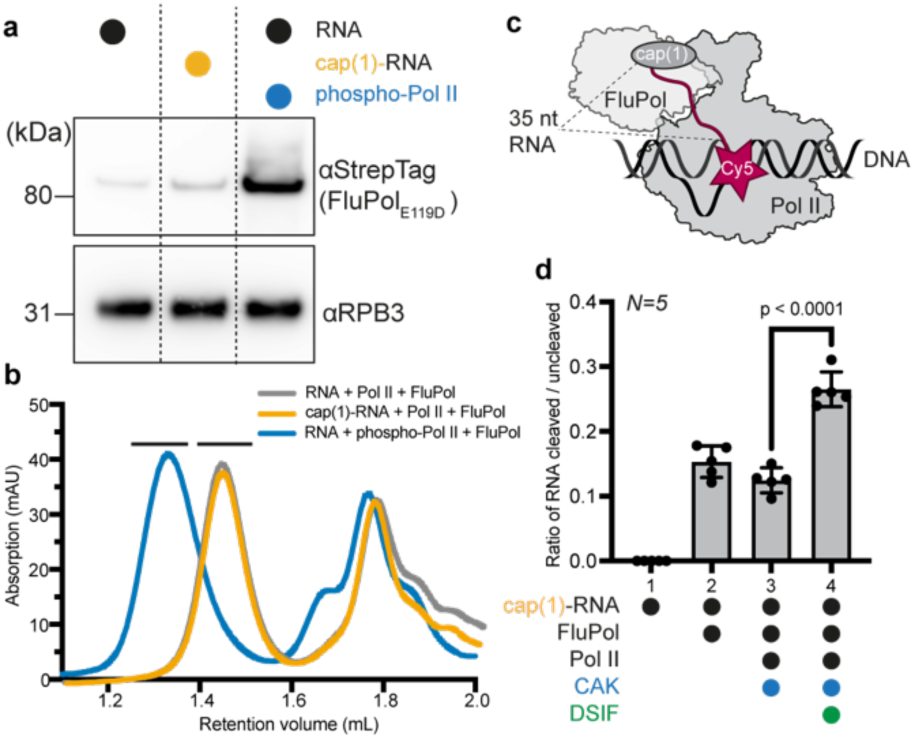
FluPol recognizes the Pol II EC. **a**, Western blot of Pol II containing peak fractions stained against RPB3 (Pol II) and Twin-Strep-Tag (FluPol subunit PB2). Different lanes represent different size exclusion chromatography runs. **b**, Absorbance at 280 nm of analytical size exclusion chromatography runs of Pol II-EC containing a 35 nt RNA with or without cap(1) and with or without CAK phosphorylation, and with FluPol. Different colors represent different chromatography runs. Black bars above the chromatogram depict Pol II complex fractions that were analyzed by Western blot in **a**. **c**, Schematic drawing of the endonuclease cleavage assay. Cap(1)-RNA is Cy5-labeled on the 3′ end. **d**, Ratio of RNA cleaved/uncleaved (intensity of cleaved product divided by intensity substrate band) in dependence of factors added. Each point reflects one experimental replicate (N=5), shown as mean ± s.d. Significance p-values were calculated using a two-tailed paired parametric t-test.

We next monitored binding of FluPol to the Pol II EC by size-exclusion chromatography (SEC) using unmodified RNA and Pol II, cap(1)-RNA or Pol II that was phosphorylated with CAK. Without CTD phosphorylation and a cap(1) structure, co-elution of FluPol with Pol II could barely be detected (**Figure 1a**). When using a cap(1)-modified RNA, the signal for FluPol in the Pol II containing peak slightly increased (**Figure 1a**). However, when the Pol II CTD was phosphorylated by CAK, the amount of FluPol associated with Pol II in the peak fractions strongly increased (**Figure 1a**). Additionally, the elution volume of the complex peak shifted towards higher molecular weight, indicating the formation of a stable complex (**Figure 1b**). Thus, the addition of a cap(1) structure to the RNA has only a modest effect on the interaction between FluPol and the Pol II EC. In contrast, phosphorylation of the Pol II CTD is the main determinant for the recruitment of FluPol to a Pol II EC, consistent with *in vivo* data demonstrating the importance of the Pol II CTD for viral transcription^2,30^.

We next asked if the increased affinity of FluPol to Pol II by CTD phosphorylation also results in enhanced endonuclease activity by FluPol. To monitor RNA cleavage, we developed a fluorescence-based assay using *in vitro*-capped RNA harboring a Cy5-label at the 3′ end (**Figure 1c**). We did not observe an increase in RNA cleavage by FluPol in the context of a phosphorylated Pol II EC compared to free RNA (**Figure 1d**, **ED Figure 1b**). This suggests that CTD phosphorylation enhances recruitment of FluPol to Pol II but the interaction of FluPol with phospho-CTD alone does not suffice to stimulate cleavage of RNA that is bound to Pol II.

Next, we tested whether the presence of the elongation factor DSIF, which binds Pol II during early elongation, stimulates the cleavage of Pol II-bound RNA. Indeed, cleavage of Pol II-bound RNA was stimulated ∼2-fold when DSIF was added to the Pol II EC in the cleavage assay (**Figure 1d**, **ED Figure 1b**). Finally, we tested whether FluPol can extend the snatched RNA primer using a radioactive FluPol RNA extension assay (Methods). We found increased FluPol-dependent RNA extension in the presence of a Pol II-DSIF EC, which is consistent with a more efficient endonuclease reaction (**ED Figure 1c**).

In summary, the cap(1) structure only has a minor impact on FluPol binding to Pol II, whereas CTD phosphorylation by CAK strongly enhances FluPol recruitment. However, CTD phosphorylation alone does not stimulate cleavage of Pol II-bound RNA by FluPol. Instead, DSIF, when added to the Pol II EC, stimulates RNA cleavage, suggesting that DSIF is part of the Pol II complex that is recognized by FluPol. Moreover, we have demonstrated that the RNA emerging from the Pol II-DSIF elongation complex can be used to prime the polymerization by FluPol. Thus, we conclude that FluPol recognizes the phosphorylated Pol II-DSIF EC as a minimal substrate for efficient cap-snatching.

### Structure of the FluPol-Pol II-DSIF cap-snatching complex

After determining the components required for efficient cap-snatching by FluPol *in vitro*, we next sought to structurally characterize a cap-snatching complex comprising FluPol, Pol II, DSIF and capped RNA by cryo-EM. To that end, we first assembled a Pol II-DSIF EC containing a 35 nt cap(1)-RNA in the presence of the CAK and ATP to phosphorylate the Pol II CTD. To capture the normally transient cap-snatching complex prior to RNA cleavage, we then added FluPol^E119D^ at low Mg^2+^ concentration, conditions in which cleavage is minimal (Methods) (**ED Figure 1d**). The complex was purified and stabilized using GraFix^41^ prior to cryo-EM sample preparation (**ED Figure 2a**). Cryo-EM data acquisition yielded 6,423,874 particles that were further sorted by 3D-classification, which yielded a subset of 369,858 particles that show good density for the Pol II-DSIF EC as well as FluPol resolved at 3.3 Å overall focused refinements of FluPol and the Pol II-DSIF EC (with respective resolutions of 2.90 Å and 2.94 Å), which allowed us to build and refine an atomic model for the complete cap-snatching complex (**Figure 2a**).

**Fig. 2.**
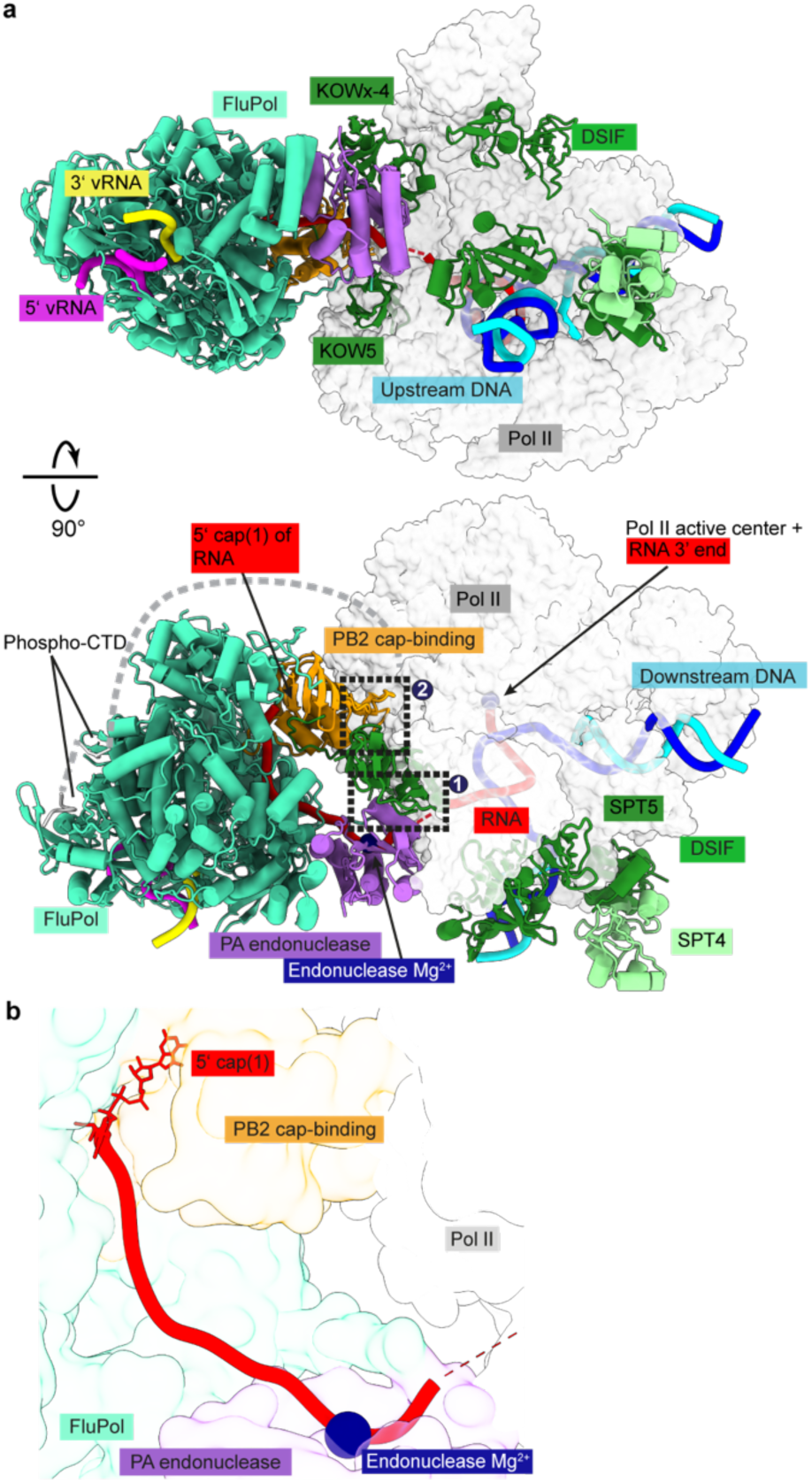
Structure of the pre-cleavage cap snatching complex. **a**, Two views of the overall structure of the pre-cleavage FluPol-Pol II-DSIF EC complex in cartoon representation except Pol II which is shown as surface. Dashed black boxes represent the locations of the two interfaces shown in **Fig. 3a-c**. The structure is shown in a FluPol side view and Pol II top view (upper half) as well as front view of FluPol and side view of Pol II. **b**, The RNA path within FluPol. Proteins are shown as transparent surfaces. The RNA is shown as ribbon tracing of the backbone. Parts of the FluPol model were removed for clarity.

The structure shows that FluPol binds to the Pol II-DSIF EC near the RNA exit channel of Pol II (**Figure 2a**). The PA endonuclease of FluPol interacts with the KOWx-4 domain of DSIF that forms a clamp around the exiting RNA in the absence of FluPol^34^ (**Figure 2a**, interface 1). In the complex, KOWx-4 is rotated ∼180° around its longitudinal axis and shifted by ∼22 Å compared to the Pol II-DSIF EC structure^34^, and the Pol II stalk containing subunits RPB4 and RPB7 is also repositioned (**ED Figure 3a,b**). The PB2 cap-binding domain of FluPol inserts in between the Pol II subunits RPB1, RPB3, and RPB11 to bind the Pol II dock domain, which is located below the RNA exit channel of Pol II (**Figure 2a**, interface 2). In line with our observation that FluPol recruitment to Pol II strongly depends on CTD phosphorylation, we observe density for phosphorylated CTD residues in two of the previously reported CTD binding sites of FluPol ^2,42,43^ (**ED Figure 3c-e**).

**Fig. 3.**
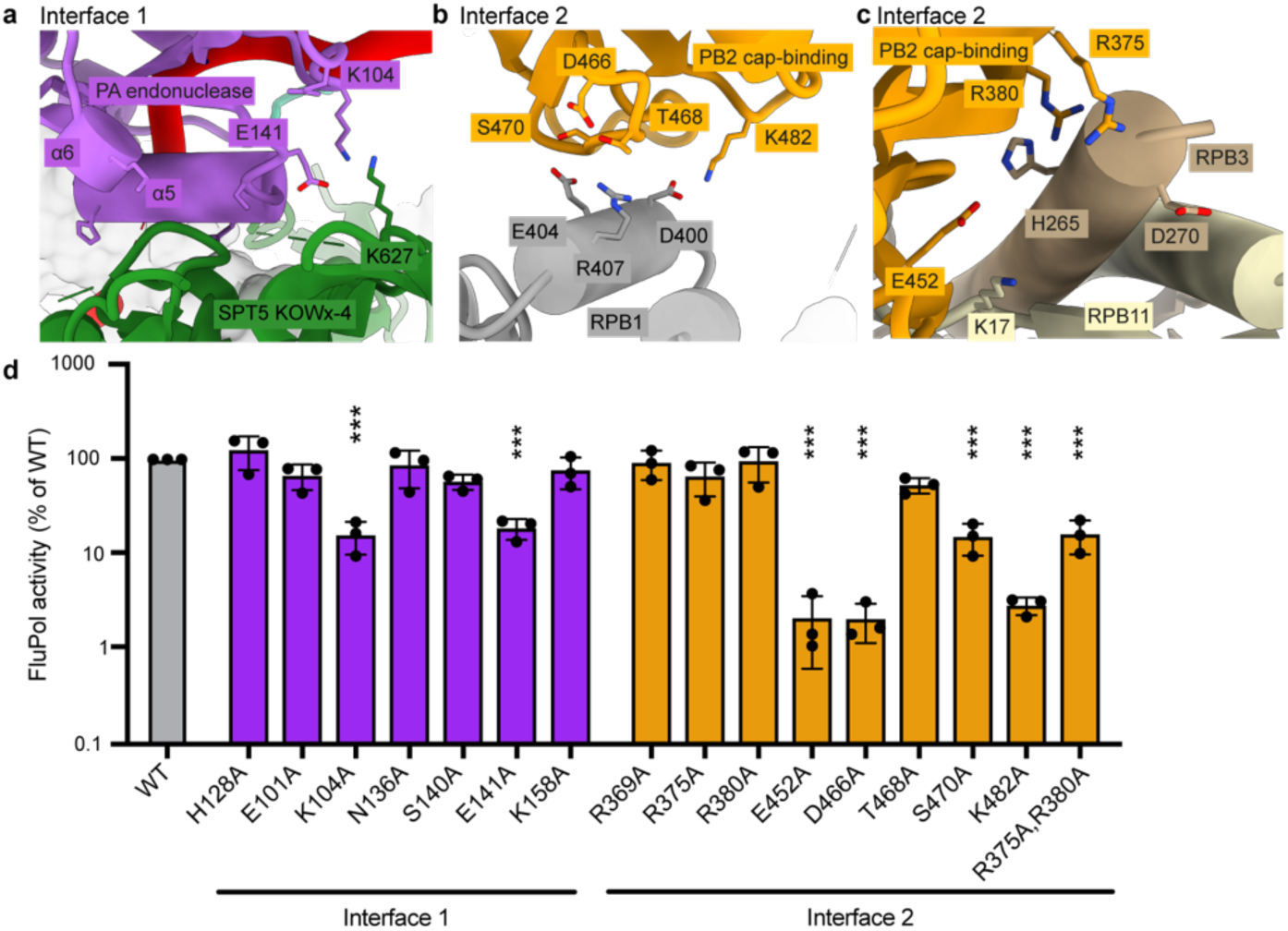
New FluPol-Pol II-DSIF EC interfaces. **a-c**, Zoom-ins on the interfaces between the FluPol PA endonuclease domain and the DSIF KOWx-4 domain (**a**), the FluPol PB2 cap-binding domain and RPB1 (**b**) or RPB3 and RPB11(**c**). Amino acids mutated are shown in stick representation and colored by heteroatoms if the mutation reduced FluPol activity significantly in a cell-based minigenome assay. Dashed black lines indicate potential interactions. **d**, Cell-based minigenome assay of A/WSN/33 FluPol activity for the indicated PA and PB2 mutants. HEK-293T cells were co-transfected with plasmids encoding PB2, PB1, PA, NP with a model vRNA encoding the Firefly luciferase. Luminescence was normalised to a transfection control and is represented as percentage of wild-type FluPol. Each point reflects one replicate (N=3), depicted as mean ± s.d., *** indicates p<0.001 as calculated by One-Way Anova - Dunnett′s multiple comparisons test referenced to wild-type.

We could trace continuous density for most of the RNA from the capped 5′-end in the PB2 cap-binding domain of FluPol all the way to the 3′-end located in the Pol II active site (**Figure 2a,b, ED Figure 3f**). This confirms that we successfully resolved the cap-snatching complex prior to endonuclease cleavage. Therefore, we called this structure the pre-cleavage complex. The cap(1) and the first four nucleotides of the RNA are well-ordered and tightly bound to the PB2 cap-binding and midlink domains as observed before^19,44^. The methylated 2′ OH of the first transcribed base packs against I260 from the PB2 midlink domain (**ED Figure 3g**), an interaction only proposed before^27^. The interaction of FluPol with the cap(1) structure is supported by parts of a previously unresolved linker between the KOWx-4 and KOW5 domains of DSIF (SPT5 residues 647-703), which interacts directly with the RNA 5′-end and the cap-binding domain of PB2 (**ED Figure 3g**). Phosphorylation of serine residues in this linker has been reported to be involved in pause release^45^.

The nucleotides between the cap-binding domain and the FluPol endonuclease could only be resolved at low resolution (**ED Figure 3f**), likely due to the flexibility of this RNA region. This precluded identification of the exact sequence register, although structural modeling (Methods) allows for 9-15 nt of RNA to be placed between the endonuclease and the cap-binding domains (**Figure 2b**), in agreement with the primer lengths of 10-15 nt that are produced by co-transcriptional cap-snatching *in vivo*^20,40^. In summary, we visualized the structure of a pre-cleavage state of FluPol bound to transcribing Pol II during cap-snatching that explains how DSIF stimulates cleavage of Pol II bound RNA.

### FluPol binding to the Pol II-DSIF EC is crucial for viral transcription *in vivo*

The biochemical analysis of FluPol endonuclease activity and the structure of the pre-cleavage complex show that FluPol binds the Pol II-DSIF EC, and that the interaction between FluPol and DSIF is important for cap-snatching *in vitro*. Next, we investigated whether the observed interactions between FluPol and the Pol II-DSIF EC are also required for FluPol activity *in vivo*, i.e. in a cellular context. For that, we used our structure of the pre-cleavage complex to identify amino acids that might be involved in the interaction between FluPol and the Pol II-DSIF EC. We chose 17 FluPol residues at the interface to the Pol II-DSIF EC that show high conservation across various influenza strains (**ED Figure 4a,b**, Methods). We then used a luciferase-based mini-genome assay to test FluPol activity in cells after mutating interface residues between PA and DSIF (**Figure 3a**) as well as PB2 and Pol II (**Figure 3b,c**) to alanine (**ED Table 2**). We focused on the FluPol variants that could be expressed at the same level as wild-type FluPol (**ED Figure 4c**).

**Fig. 4.**
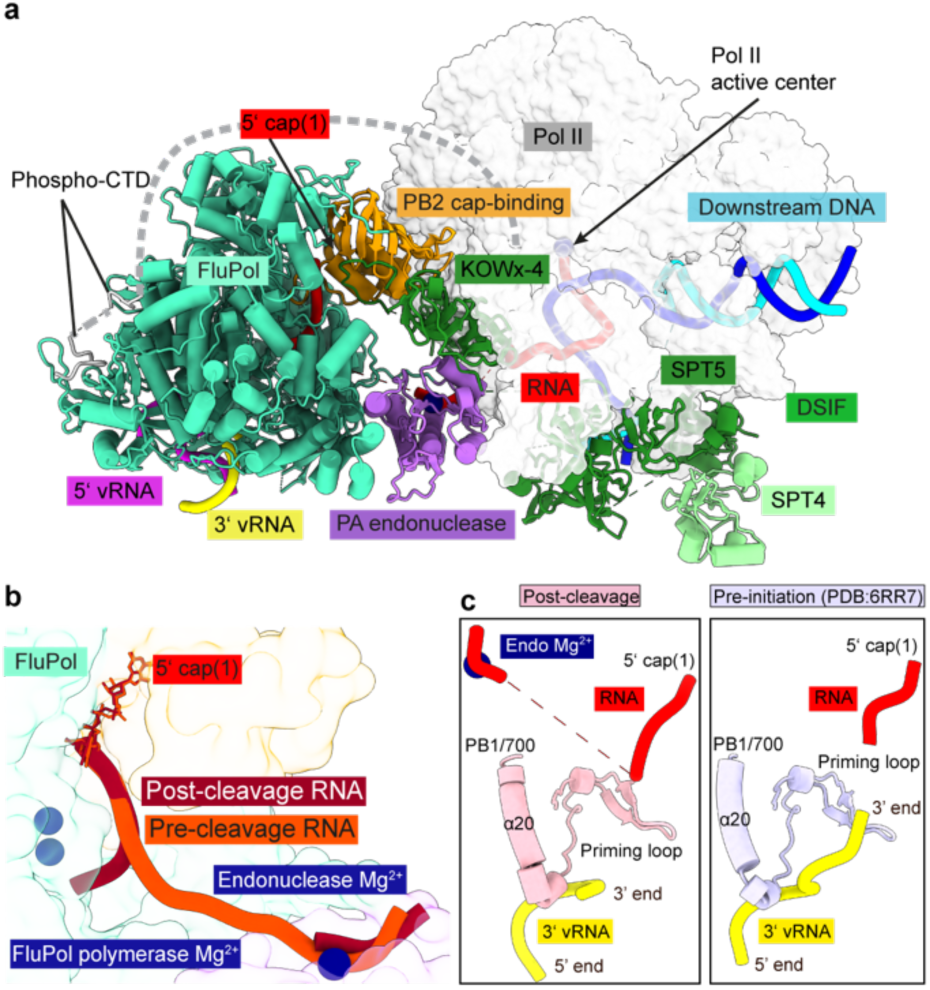
Structure of the post-cleavage cap snatching complex. **a**, Overall structure of the post-cleavage FluPol-Pol II-DISF EC complex in cartoon-style representation except Pol II which is shown as surface. **b**, Comparison of the RNA-path in FluPol between pre and post-cleavage complex. Proteins are shown as transparent surfaces and the RNA is shown as ribbon tracing of the backbone. FluPol polymerase active site Mg^2+^ atoms are modeled based on the FluPol elongation complex^15^. Parts of FluPol were removed for clarity. **c**, Comparison of the FluPol polymerase active site conformations in the post-cleavage (pink) and pre-initiation (PDB:6RR7, light purple^44^) states. Only the priming loop, the viral mRNA the 3′ vRNA are shown.

When we mutated PA residues involved in the interaction with DSIF (**Figure 3a**), mutations in the α5 and α6 helices of PA (H128, E101, N136, S140 and K158, respectively) did not reduce viral transcription *in vivo* (**Figure 3d**). However, mutating K104 or E141 to alanine reduced FluPol activity ten-fold in the mini-genome assay (**Figure 3d**). Both residues are close to the conserved DSIF residue K627, with which PA E141 is likely involved in forming a salt bridge that might also be stabilized by the nearby K104 (**Figure 3a, ED Figure 4d**). Thus, we show that the integrity of the PA endonuclease interface with DSIF is vital for efficient FluPol activity *in vivo*. This also agrees with our biochemical data showing that DSIF stimulates cleavage of Pol II-bound RNA by FluPol *in vitro* (**Figure 1d**).

On the surface of PB2, we mutated residues involved in interacting with the Pol II subunits RPB1, RPB3 and RPB11 (**Figure 3b,c**). PB2 residues D466, T468, S470 and K482 are located at the interface between the PB2 cap-binding domain and the RPB1 dock domain and can form hydrogen bonds as well as salt bridges with RPB1 residues E400, D404, and R407, respectively. From this interface, the T468A mutant retained ∼60% of wild-type activity, whereas all other mutations almost abolished FluPol activity (**Figure 3d**). Mutation of PB2 E452, which might form a salt bridge with K17 of RPB11, also reduced FluPol activity *in vivo* (**Figure 3d**). While the individual mutations of PB2 residues R375 and R380 that might interact with RPB3 do not significantly reduce activity, they did when mutated together (**Figure 3d**). Additionally, the interface residues in RPB1, RPB3 and RPB11 are highly conserved between mammals and birds (**ED Figure 4e-g**). These results show that the interface between the PB2 cap-binding domain and the Pol II surface is important for FluPol activity *in vivo*.

We conclude that both interfaces between PA and DSIF, as well as PB2 and the Pol II surface, are crucial for efficient cap-snatching and viral transcription *in vivo*. Since we observed perturbations of viral transcription by mutating residues that are conserved across several influenza strains, we propose that FluPol of other influenza A strains binds to the Pol II-DSIF EC in a similar way to that observed in the structure of the pre-cleavage complex. Thus, we establish the molecular interfaces between FluPol and the Pol II-DSIF EC during co-transcriptional cap-snatching in cells.

The pre-cleavage complex structure reveals the RNA trajectory direct from the cap-binding to the endonuclease domain of FluPol, which is clearly incompatible with the FluPol pre-initiation complex that precedes viral transcription^19,44^. This suggests that FluPol, the primer RNA or both must undergo conformational changes after endonuclease RNA cleavage to position the newly generated RNA 3′-end in the FluPol PB1 polymerase active site for RNA extension. To investigate these structural transitions, we resolved a cap-snatching complex of FluPol bound to the Pol II-DSIF EC after the PA endonuclease has cleaved the RNA.

To achieve that, we assembled the Pol II-DSIF EC with cap(1)-RNA and FluPol^E119D^ as before, but in the presence of 3 mM Mg^2+^ (**ED Figure 5a**), which allowed for RNA cleavage during cryo-EM sample preparation (**ED Figure 1d)**. We then performed cryo-EM as for the pre-cleavage complex (**Figure 4a**, **ED Figure 5b-e, ED Figure 6**). The resulting post-cleavage structure is very similar to the pre-cleavage complex (**ED Figure 7a**), except for differences in the primer RNA. In particular, we could only trace the RNA from the Pol II active site until the PA endonuclease active site, after which the density discontinues abruptly (**ED Figure 7b**). The 5′ cap(1) structure remains bound as before in the cap-binding site. However, the cleaved RNA 3′ end points towards the FluPol polymerase active site (**Figure 4b**). The endonuclease cleaves a fragment of ∼10-15 nt from the Pol II transcript^20,40^, of which we can observe cryo-EM density for the first 7 nt of the primer in the post-cleavage complex, indicating that the missing nucleotides are disordered. Furthermore, in the post-cleavage complex, the priming loop near the FluPol polymerase active site is still extended and ordered (**Figure 4c, ED Figure 7c**), as expected when the snatched RNA primer has not yet base paired with a viral RNA template 3′ end. Thus, FluPol in the post-cleavage cap-snatching state resembles the FluPol pre-initiation complex that was previously reported^19,46^ (**ED Figure 7d**).

**Fig. 5.**
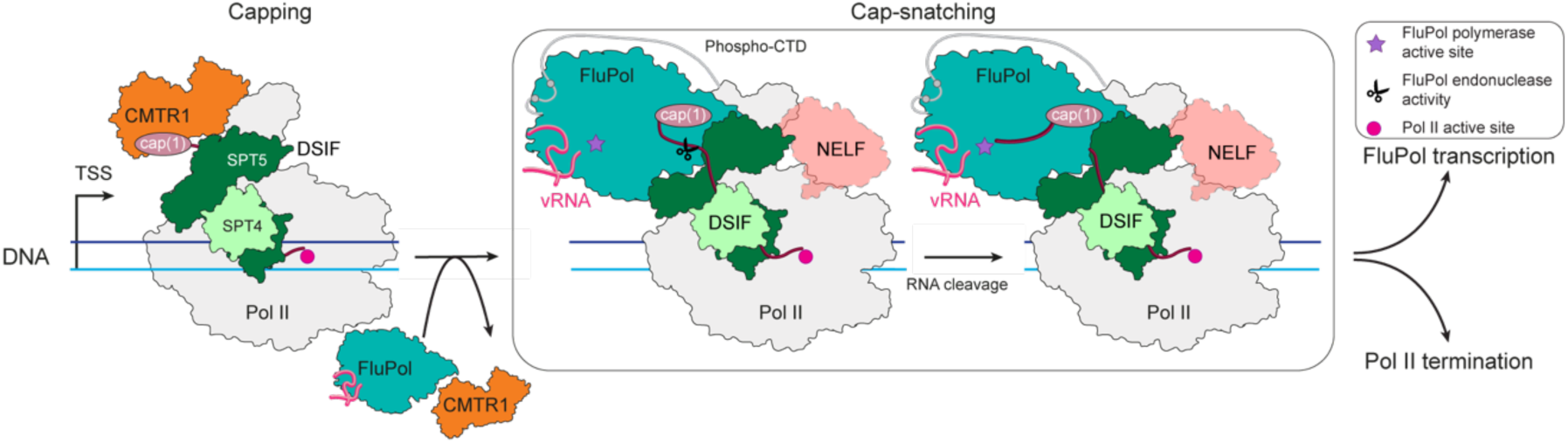
Model of co-transcriptional cap-snatching. Capping enzymes, including CMTR1, synthesize the cap(1) structure on the RNA co-transcriptionally. After capping is finished, CMTR1 dissociates from Pol II. The resulting Pol II-DSIF EC with a capped RNA is a substrate for cap-snatching and is bound by FluPol. Then, the FluPol endonuclease cleaves the RNA, FluPol may dissociates from the Pol II EC surface and initiates transcription, whereas Pol II gets terminated.

In summary, RNA cleavage by the FluPol endonuclease leads to rearrangements of the capped RNA primer and a new RNA trajectory that is indicative of a FluPol pre-initiation complex. Thus, the PA endonuclease activity on Pol II-bound capped RNA leads to a state of FluPol that is ready to initiate viral transcription with minimal conformational changes. Therefore, a rotation of the cap-binding domain does not seem to be required to direct the cleaved primer into the polymerase active site as previously proposed^47^.

## Discussion

The results presented here close a major gap in our understanding of the life cycle of one of the most common human viral pathogens. By combining structural, biochemical and cellular approaches, we propose a molecular mechanism of cap-snatching by FluPol, which involves three major steps (**Figure 5, ED Movie 1**). First, FluPol directly binds to the host transcription machinery. The minimal substrate for efficient cap-snatching is a Pol II-DSIF EC with a cap(1)-RNA and phosphorylated Pol II CTD, which is found during early host transcription^24,25,34^. Second, RNA cleavage by the FluPol endonuclease generates a 10-15 nt primer. After cleavage, the new 3′end of the capped RNA primer swings towards the FluPol polymerase active site, resulting in a conformation similar to a FluPol pre-initiation complex. Third, the 3′ end of the capped RNA primer can anneal to the vRNA template, and FluPol elongates the viral mRNA.

Building on our mechanistic understanding of co-transcriptional cap-snatching, our model also provides insights into how cap-snatching is coordinated with host transcription by Pol II (**Figure 5**). Early Pol II transcription includes RNA capping, early elongation, promoter-proximal pausing, pause release, and premature termination, which have been structurally characterized^26,48–50^. The comparison of the cap-snatching complex structure with CMTR1 bound to the Pol II-DSIF EC^26^ shows that binding of FluPol and CMTR1 to Pol II is likely mutually exclusive (**ED Figure 8a**). Thus, for cap-snatching to occur, CMTR1 likely has to dissociate after addition of the essential 2′-OH methylation to the RNA cap^27^. CMTR1 dissociation from the Pol II-DSIF EC then allows for FluPol binding to the RNA and the Pol II-DSIF surface as observed in the pre-cleavage structure (**Figure 2a**). NELF binding to the Pol II-DSIF EC establishes the PEC^48^, which can accommodate FluPol binding without clashes (**ED Figure 8b**). Active elongation in the EC* and termination factors such as Integrator or XRN2, however, clash with FluPol when bound to Pol II at the same time (**ED Figure 8c-e**). Thus, the window of opportunity for cap-snatching likely opens during Pol II early elongation (Pol II-DSIF EC) and pausing (PEC), and closes upon pause release and formation of the EC*. Within that window of opportunity, early elongating and paused Pol II represent relatively long-lived substrates for cap-snatching as Pol II resides in this phase up to several minutes^51,52^. FluPol binding to the KOWx-4-KOW5 linker of SPT5 might further extend the residence time of Pol II in the paused state by preventing phosphorylation of the linker, which was shown to be important for pause release^45^. Although the exact fate of FluPol after cap-snatching remains enigmatic, FluPol might remain bound on the Pol II surface during the very first steps of viral transcription elongation (**ED Figure 7e**)^36^. However, as FluPol transcription proceeds, the elongating viral mRNA will extrude more and more out of the FluPol product exit channel, requiring more space to be accommodated. In addition, FluPol undergoes conformational changes during the initiation to elongation transition^15,19^. These events might lead to FluPol dissociation from the Pol II core (although it could remain bound to the CTD^42^), perhaps concomitant with release of the capped RNA from the FluPol cap-binding site and subsequent recruitment of the nuclear cap-binding complex^53^. Alternatively, Integrator or XRN2 binding to the Pol II surface could compete with FluPol, triggering its dissociation (**ED Figure 8d,e**).

Our structures of co-transcriptional cap-snatching complexes have established the molecular template for the search of inhibitors that may disrupt conserved interfaces crucial for cap-snatching. Targeting of such small protein-protein interfaces is inherently difficult^54^. However, together with recent advances in predicting such interactions^55,56^, our structure may catalyze future *in silico* and experimental studies to identify compounds suitable to disrupt the interface between FluPol and transcribing Pol II.

## Acknowledgments

We thank R. Muir and N. Iwicki for help with purifying FluPol variants, F. Grabbe for providing purified CAK, U. Steuerwald for support at the electron microscope and maintaining the cryo-EM facility, T. Schulz for providing pig thymus tissue, P. Rus and U. Neef for running the insect cell facility and J. Walshe and M. Ochmann for advice on data processing, S. Vos, C. Bernecky and T. Kouba for help with preliminary experiments and S. Paisant for help with plasmid mutagenesis.

## Author Contributions

A.H.R., M.Li., M.Lu., C.D., P.C., N.N., and S.C. designed the study. A.H.R. and D.L. planned all experiments except for the cell-based minigenome assays. T.K. and N.N. planned cell-based assays. A.H.R., D.L., C.O., A.S. prepared protein components. D.L. performed biochemical assays and prepared samples for cryo-EM. A.H.R., I.F. and C.D. acquired and analyzed cryo-EM data. A.H.R. and S.C. built the molecular models. M.Lu. pioneered reconstitution of the cap-snatching complex ^31^, T.K. and M.D. performed cell-based assays. D.L. and A.H.R. designed figures. A.H.R. and C.D. wrote the manuscript with input from all authors. All authors read and approved the final manuscript. Conceptualization: P.C., S.C.; Methodology: A.H.R., D.L., S.C.; Formal analysis: A.H.R., D.L., I.F., M.Li., T.K., N.N.; Investigation: A.H.R., D.L., M.D., T.K., M.Lu.; Writing - original draft preparation: A.H.R.; Visualization: D.L., A.H.R.; Writing - review and editing: A.H.R., D.L., T.K., M.Li., M.Lu. C.D., P.C., N.N., and S.C; Funding acquisition: P.C., A.H.R., N.N.; Resources: A.H.R., D.L., C.O., A.S.; Supervision: P.C., M.Li., C.D., S.C., N.N.

## Data Availability

The electron density reconstructions and final models were deposited with the EM Data Bank (accession codes 50892, and 50927) and the PDB (accession codes PDB 9FYX, and 9G0A).

## Extended Data Figures

**Extended Data Fig. 1.**
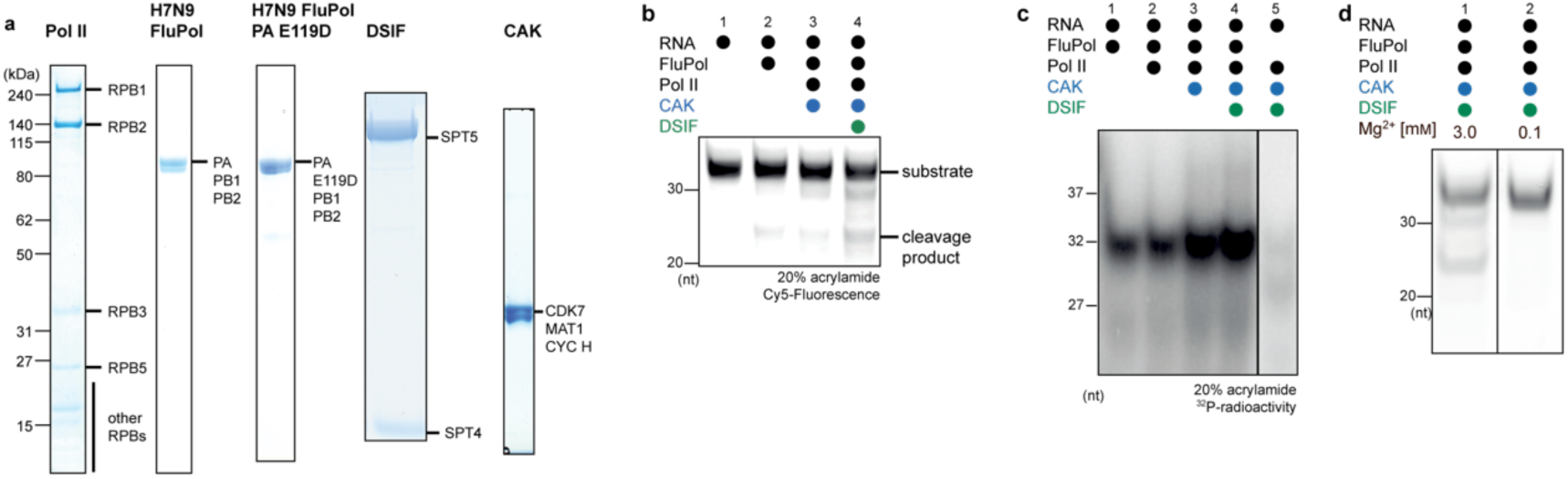
Related to Fig. 1. FluPol recognizes the Pol II EC. **a,** Representative images of Coomassie-stained SDS-PAGE lanes of the purification of the single protein components used. Protein band assignments are based on size. **b**, Representative example for the denaturing urea PAGE of an endonuclease assay. The substrate and the product bands are labeled. **c**, Scintillation image of denaturing PAGE radioactivity elongation assay showing increased FluPol transcription upon DSIF addition and CTD phosphorylation. **d**, Denaturing urea PAGE of an endonuclease assay with varying Mg^2+^ concentrations.

**Extended Data Fig. 2.**
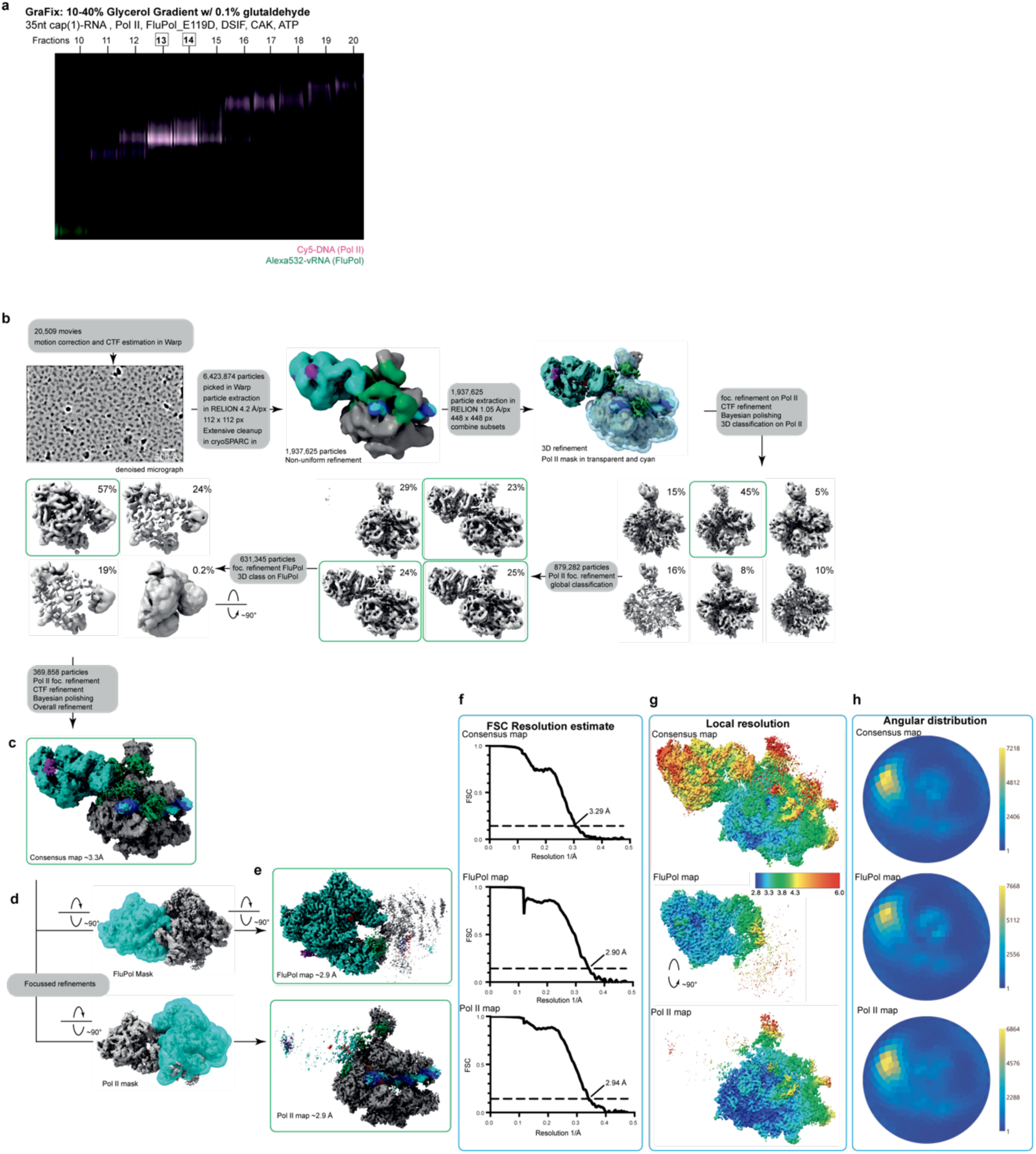
Data acquisition and processing of the pre-cleavage complex. **a**, Fluorescence scan of the native PAGEs analyzing GraFix gradient fractions of the pre-cleavage complex preparation. The image is an overlay of Cy5 and ATTO532 signals. In magenta is the scan for the Cy5 channel of the labeled template DNA, and in green is the scan for the ATTO532 channel of the labeled 3′ vRNA. Highlighted fractions were combined and used for cryo-EM analysis. **b,** Flowchart illustrating the key steps of the processing pipeline for obtaining the structure of the pre-cleavage complex. **c,** consensus density map of the pre-cleavage FluPol-Pol II-DSIF EC complex colored by underlying protein components. **d,e,** Masks used for focused refinements and their resulting maps. **f,** Gold-Standard Fourier shell correlation plots of the consensus and focused maps. **g,** Local resolution of consensus and focus refinement maps. Local resolution estimations were performed in RELION. **h**, Angular distribution of the

**Extended Data Fig. 3.**
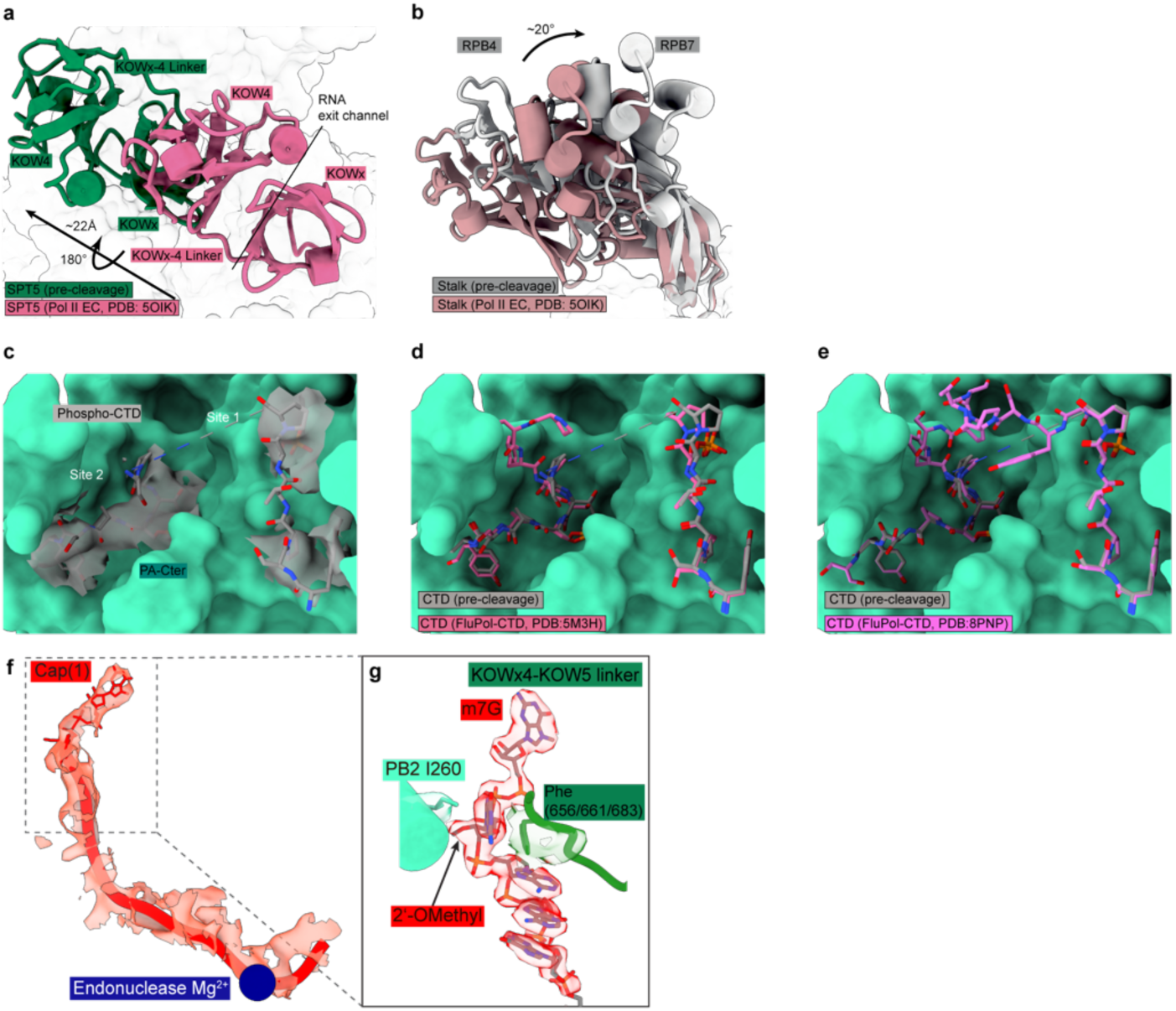
Related to Fig. 2. Structure of the pre-cleavage cap snatching complex. **a-b,** Comparison of pre-cleavage structure with the canonical Pol II-DSIF EC (PDB:5OIK)^34^. **a**, The canonical position of the KOWx-4 domain is shown in pink, and the position observed in the pre-cleavage structure is shown in green. The KOWx-4 domain of DSIF SPT5 is displaced by ∼22 Å and rotated by ∼180° relative to the canonical conformation. **b**, The canonical position of the Pol II stalk is colored in dusty pink, and the stalk model from the pre-cleavage structure is depicted in gray. The Pol II stalk is moved by 20° around its base relative to the canonical conformation. **c,** Phosphorylated CTD of RPB1 bound to FluPol shown in sticks. FluPol is shown as surface. The obtained cryo-EM density is displayed in transparent gray. **d,e**, Comparison of CTD binding the prior structures ^2,42^ shows a similar conformation of the CTD in the CTD-binding site of FluPol. **f**, Cryo-EM density for the RNA within FluPol between the PB2 cap-binding and PA endonuclease domains. **g**, 5′ mRNA cap(1) inside the PB2 cap-binding domain and the supporting Phe from the KOWx-4-KOW5 linker. The density for the RNA and the linker is transparent and colored in the color of the underlying models.

**Extended Data Fig. 4.**
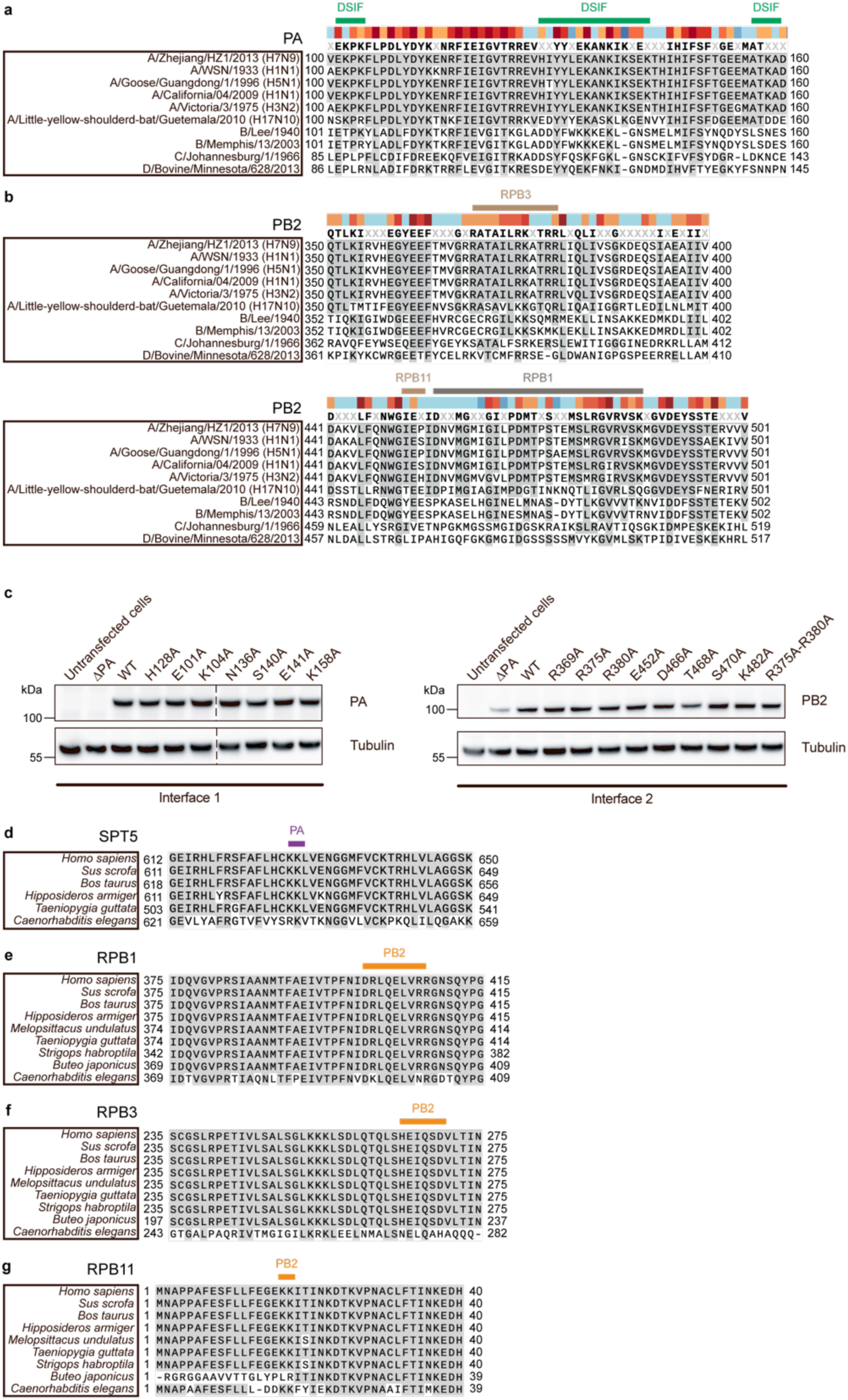
Evolutionary conservation of FluPol and Pol II residues involved in the interface. **a**, Sequence alignment of PA from representative influenza A–D strains. Residue stretches interacting with DSIF are highlighted with a green box above the alignment. **b,** Sequence alignment of PB2 from representative influenza A–D strains. Residue stretches interacting with different RPBs are highlighted with boxes in different colors above the alignment. **c**, Western blots against PA, PB2, and tubulin for wild type and mutant FluPol transiently expressed in HEK-293T cells. **d-g**, Sequence alignments of SPT5 (**d**), RPB1 (**e**), RPB3 (**f**), RPB11 (**g**) from mammals (*H. sapiens, S. scrofa, B. taurus, H. armiger*), birds (*M. undulatus,* FluPol are highlighted with a box above the alignment. Bird species selection was based on well annotated RPB1. SPT5 was not well annotated in *M. undulatus, S. habroptila* and *B. japonicus*, consequently, they were omitted in panel **d**.

**Extended Data Fig. 5.**
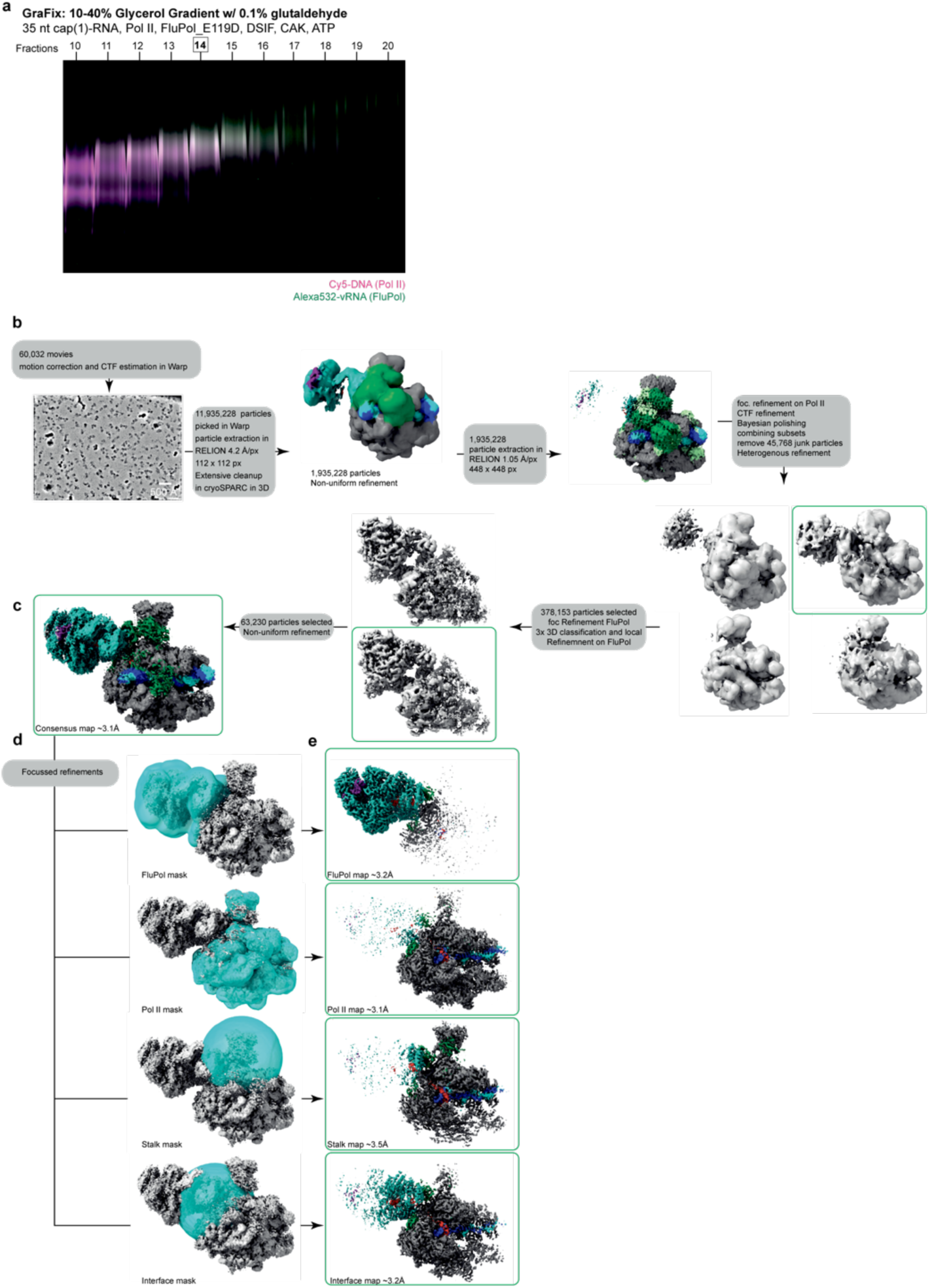
Data acquisition and processing of the post-cleavage complex. **a**, Fluorescence scan of the native PAGEs analyzing GraFix gradient fractions of the post-cleavage complex preparation. The image is an overlay of Cy5 and ATTO532 signals. In magenta is the scan for the Cy5 channel of the labeled template DNA, and in green is the scan for the ATTO532 channel of the labeled 3′ vRNA. Highlighted fractions were combined and used for cryo-EM analysis. **b,** Flowchart illustrating the key steps of the processing pipeline for obtaining the post-cleavage FluPol-Pol II-DSIF EC structure. **c,** Consensus density map of the post-cleavage FluPol-Pol II-DSIF EC colored by underlying protein components. **d,e,** Masks

**Extended Data Fig. 6.**
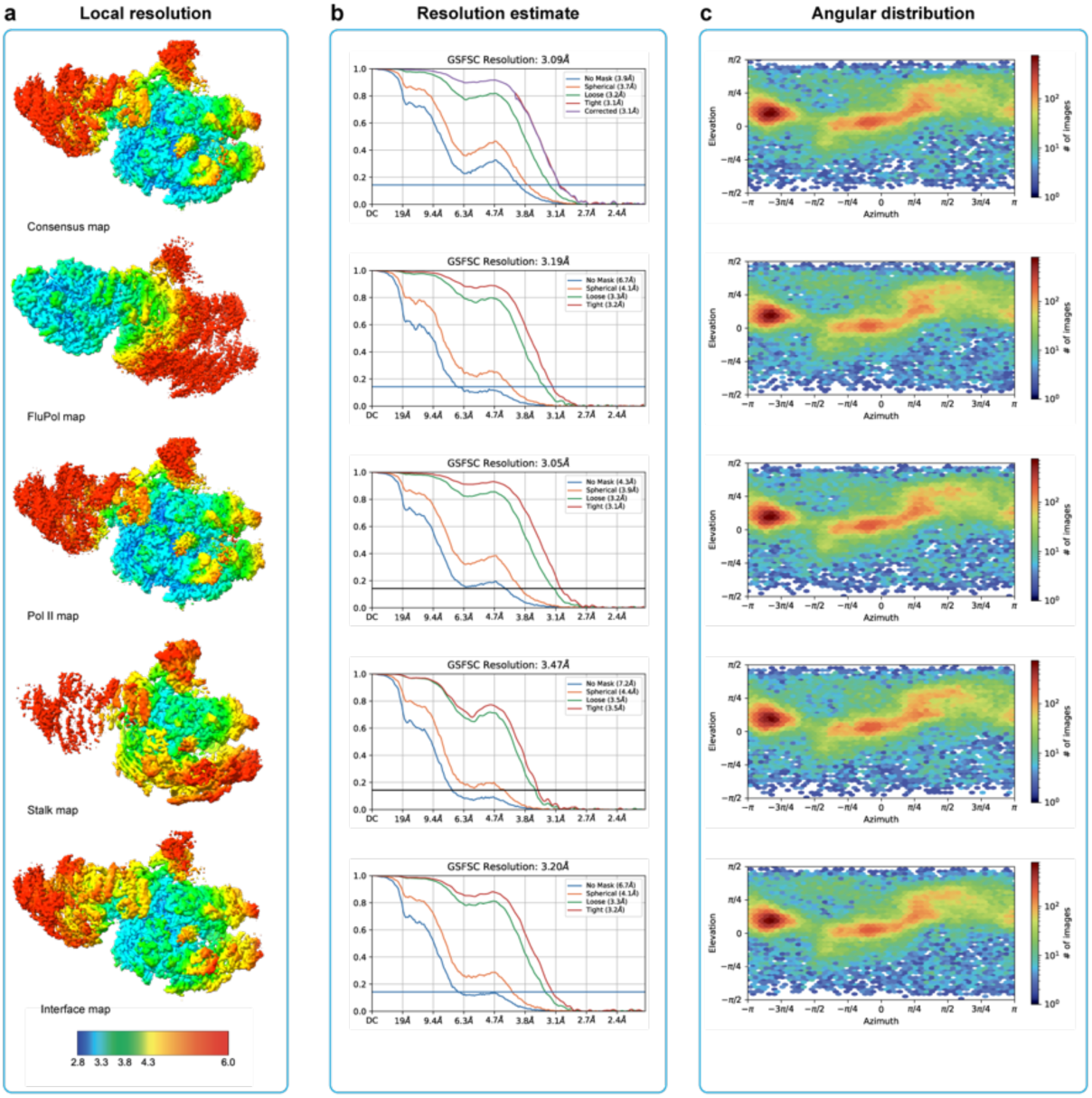
Data quality of the post-cleavage structure cryo-EM data. **a**, Local resolution of consensus and focus refinement maps. Local resolution estimations were performed in RELION. **b,** Gold-standard Fourier shell correlation plots of the consensus and focused maps. **c**, Angular distribution of the consensus refinement and focused refinements as plotted by cryoSPARC.

**Extended Data Fig. 7.**
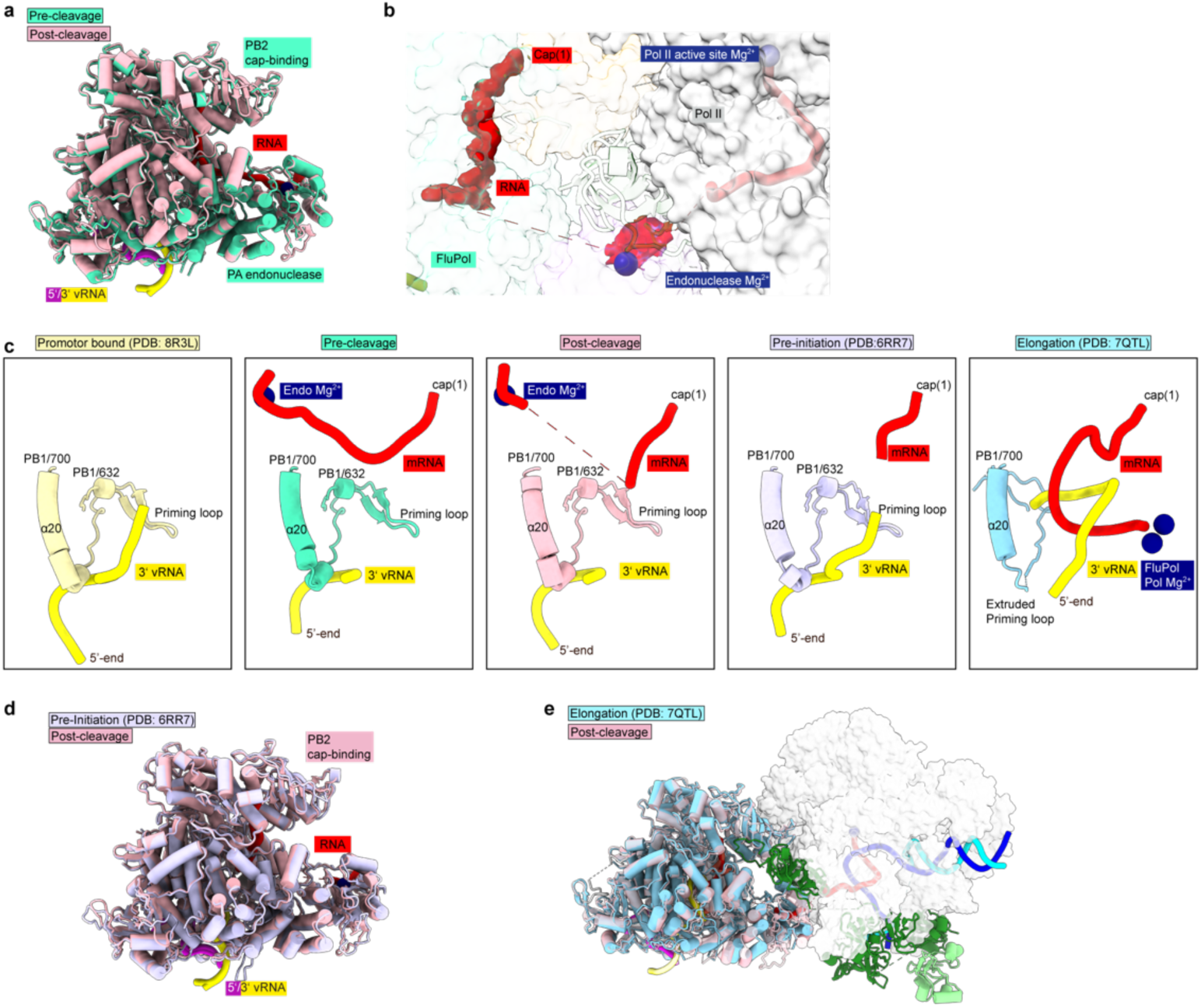
Comparison of FluPol structures. **a**, Comparison of pre- and post-cleavage FluPol structures. Pre-cleavage is shown in turquoise, and post-cleavage is pink. **b,** RNA-path within FluPol. Proteins are shown as transparent surfaces. The RNA is shown as ribbon together with the FluPol map. The density is colored by the underlying model part and restricted by the distance to the RNA in ChimeraX. **c**, Comparison of FluPol transcription pre-initiation state (PDB: 4WSB, yellow)^47^ with the pre-cleavage cap-snatching state (turquoise), post-cleavage cap-snatching state (pink), pre-initiation state (PDB: 6RR7, light purple)^44^ and elongation state (PDB: 7QTL, light blue)^36^. Only the priming loop, the 3′ vRNA, and the capped RNA are depicted. **d**, Comparison of post-cleavage and pre-initiation FluPol structure. Post-cleavage is shown in pink, and pre-initiation in transparent white (PDB:6RR7)^44^. **e**, superposition of an early elongating FluPol onto the post-cleavage FluPol-Pol II-DSIF EC reveals no clash between the early elongating FluPol and the Pol II EC (PDB: 7QTL, light blue)^36^.

**Extended Data Fig. 8.**
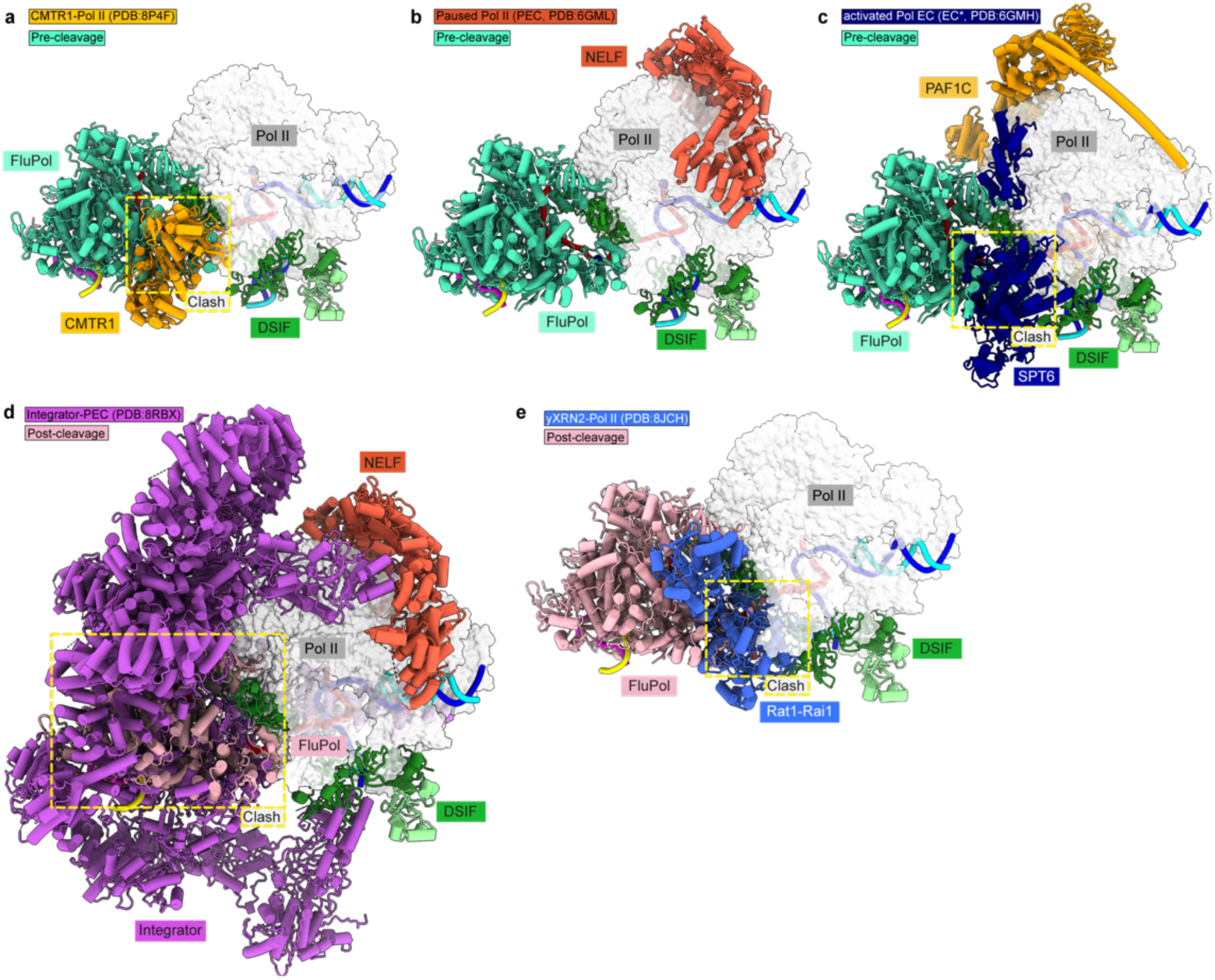
Cap-snatching is compatible with early Pol II elongation and pausing. **a**, Overlay of the pre-cleavage complex (this study) and the Pol II-CMTR1 EC (PDB:8P4F) shows that FluPol clashes with CMTR1^26^. **b**, NELF binding to Pol II is compatible with the cap-snatching as NELF and FluPol bind to different regions of Pol II (PDB: 6GML)^48^. **c**, FluPol clashes with SPT6 core in the activated elongation complex (PDB: 6GMH)^49^. **d,** Comparison of the post-cleavage structure with the Integrator complex bound to the paused Pol II EC (PDB: 8RBX)^50^ shows a clash between FluPol and the Integrator cleavage module. **e**, Comparison of the post-cleavage structure with yeast Rat1-Rai1 (homolog of human XRN2) bound to a yeast Pol II EC (PDB: 8JCH)^57^ shows a clash between FluPol and Rat1-Rai1.

**Extended Data Table 1:** Cryo-EM data acquisition, processing, and refinement statistics.

|  | Pre-cleavage<br>(EMD-50892, PDB 9FYX) | Post-cleavage<br>(EMD-50927, PDB 9G0A) |
| --- | --- | --- |
| **Data collection and processing** |  |  |
| Magnification | 81,000 | 81,000 |
| Voltage (kV) | 300 | 300 |
| Electron exposure (e-/Å<sup>2</sup>) | 39.94 | 40.0 |
| Defocus range (µm) | -0.5 to -2.0 | -0.5 to -2.0 |
| Pixel size (Å) | 1.05 | 1.05 |
| Symmetry imposed | C1 | C1 |
| Initial particle images (no.) | 6,423,874 | 11,935,228 |
| Final particle images (no.) | 369,858 | 63,230 |
| Map resolution (Å) | 3.3 | 3.1 |
| FSC threshold | 0.143 | 0.143 |
| Map resolution range (Å) | 2.74-11.4 | 2.85-11.9 |
| Map sharpening *B* factor (Å<sup>2</sup>) | -72.3 | -60.7 |
| **Refinement** |  |  |
| Initial model (PDB) | 7B0Y, 5OIK,7QTL | 7B0Y, 5OIK,7QTL,7YCX |
| Model resolution (Å) | 3.3 | 3.1 |
| **Model composition** |  |  |
| Protein residues | 6690 | 6699 |
| Nucleic acid residues | 122 | 106 |
| Ligands | GGG:1, Zn:9, Mg:2, PO4:2 | GGG:1, Zn:8, Mg:2, PO4:3 |
| ***B* factors (Å<sup>2</sup>)** |  |  |
| Protein | 98.79 | 109.27 |
| Nucleotides | 1117.71 | 149.9 |
| Ligand | 61.41 | 144.58 |
| **r.m.s. deviations** |  |  |
| Bond lengths (Å) | 0.004 | 0.008 |
| Bond angles (°) | 0.686 | 1.168 |
| **Validation** |  |  |
| MolProbity score | 1.64 | 1.86 |
| Clashscore | 7.60 | 12.97 |
| Poor rotamers (%) | 0.10 | 0.08 |
| **Ramachandran plot** |  |  |
| Favored (%) | 96.54 | 96.39 |
| Allowed (%) | 3.46 | 3.61 |
| Disallowed (%) | 0.00 | 0.00 |

**Extended Data Table 2:**
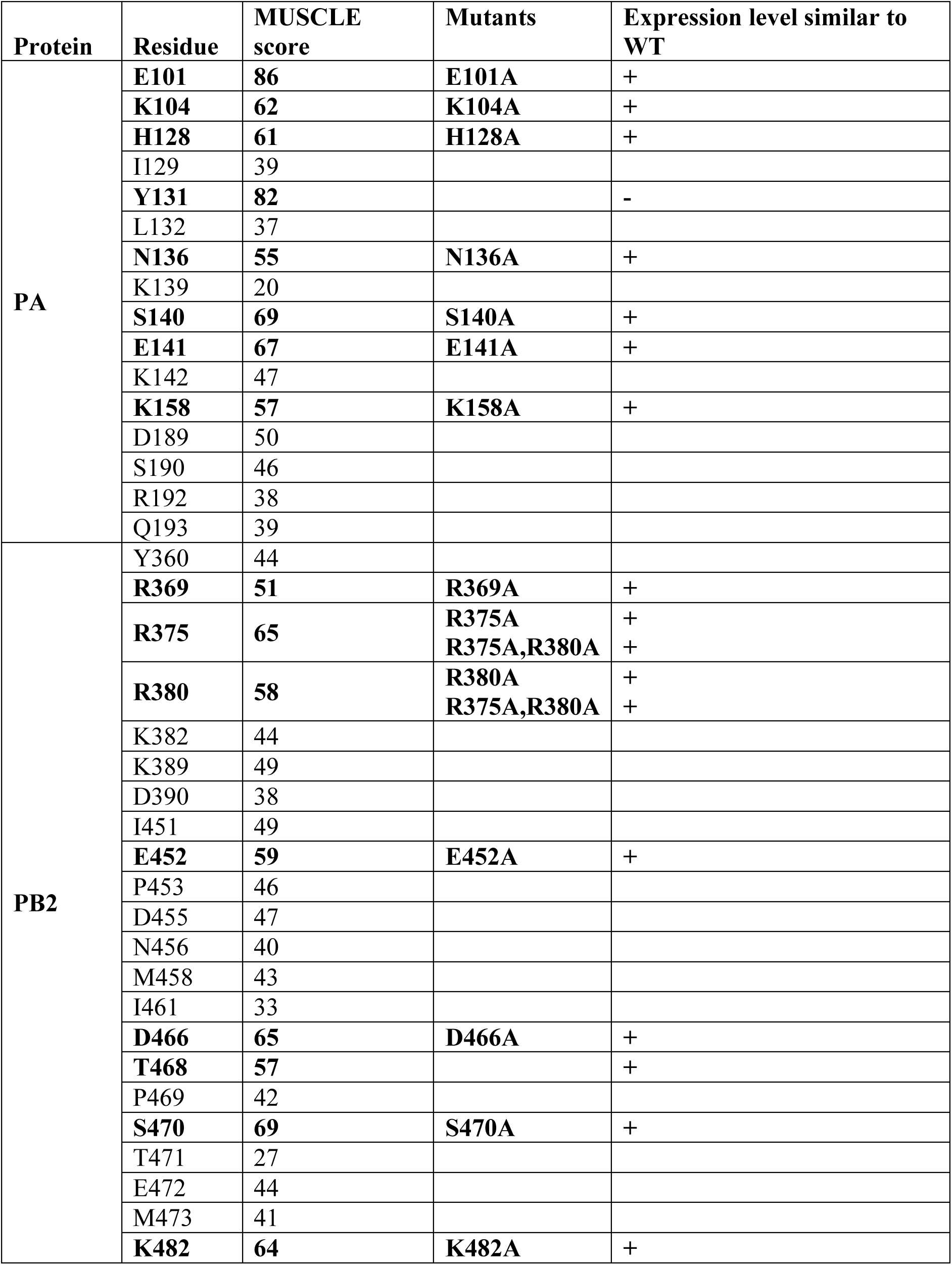
Surface Residues, their conservation score. Conserved residues are printed in bold. For conserved residues, noted mutants are checked for the expression level.

| Protein | Residue | MUSCLE score | Mutants | Expression level similar to WT |
| --- | --- | --- | --- | --- |
| PA | **E101** | **86** | **E101A** | + |
|  | **K104** | **62** | **K104A** | + |
|  | **H128** | **61** | **H128A** | + |
|  | I129 | 39 |  |  |
|  | **Y131** | **82** |  | - |
|  | L132 | 37 |  |  |
|  | **N136** | **55** | **N136A** | + |
|  | K139 | 20 |  |  |
|  | **S140** | **69** | **S140A** | + |
|  | **E141** | **67** | **E141A** | + |
|  | K142 | 47 |  |  |
|  | **K158** | **57** | **K158A** | + |
|  | D189 | 50 |  |  |
|  | S190 | 46 |  |  |
|  | R192 | 38 |  |  |
|  | Q193 | 39 |  |  |
| PB2 | Y360 | 44 |  |  |
|  | **R369** | **51** | **R369A** | + |
|  | **R375** | **65** | **R375A** | + |
|  |  |  | **R375A,R380A** | + |
|  | **R380** | **58** | **R380A** | + |
|  |  |  | **R375A,R380A** | + |
|  | K382 | 44 |  |  |
|  | K389 | 49 |  |  |
|  | D390 | 38 |  |  |
|  | I451 | 49 |  |  |
|  | **E452** | **59** | **E452A** | + |
|  | P453 | 46 |  |  |
|  | D455 | 47 |  |  |
|  | N456 | 40 |  |  |
|  | M458 | 43 |  |  |
|  | I461 | 33 |  |  |
|  | **D466** | **65** | **D466A** | + |
|  | **T468** | **57** |  | + |
|  | P469 | 42 |  |  |
|  | **S470** | **69** | **S470A** | + |
|  | T471 | 27 |  |  |
|  | E472 | 44 |  |  |
|  | M473 | 41 |  |  |
|  | **K482** | **64** | **K482A** | + |

## Material and Methods

### Cloning and purification of proteins

To generate H7N9 FluPol with impaired endonuclease activity, the PA E119D mutation was introduced into the PA gene. A pFastBac Dual vector encoding the influenza polymerase heterotrimer subunits of A/Zhejiang/DTID-ZJU01/2013 (H7N9)^36^, was used as a template for PCR site-directed mutagenesis and Gibson cloning. Sequencing of the polymerase subunits confirmed the successful introduction of the E119D mutation in the PA gene.

The wild-type FluPol and FluPol PA E119D were essentially expressed and purified as described in ^36^ with the following modification for all experiments, except the sample preparation for the post-cleavage structure. Instead of ammonium sulfate precipitation, the supernatant was clarified by ultracentrifugation in a Ti45 rotor (Beckman Coulter) at 45.000 rpm and 4°C for 1 h.

The human transcription factors (DSIF and CAK kinase trimer) were expressed and purified as described previously^26,34,58^. Pol II was purified from pig thymus as described in ^34,48^, leaving out the size exclusion step.

### *In vitro* transcription

The mRNAs were transcribed from two DNA primers^59^. The primers are complementary at the promoter site for the T7 polymerase, and the desired RNA sequence is single-stranded. The *in vitro* transcription mixture contained 1 µM primers, 40 mM Tris-HCl pH 8.0, 30 mM MgCl_2_, 2 mM spermidine, 50 mM NaCl, 5 mM NTPs (pH adjusted to 7), 2% DMSO, 0.01% TritonX-100, and 5% T7 DNA-dependent RNA polymerase (homemade). The *in vitro* transcription reaction was incubated at 37°C overnight.

The following day, for 1 mL of reaction, 10 µL of Proteinase K (NEB) and 10 µL of DNAse I (ThermoFisher) were added. The reaction was incubated at 37°C for another 10 min. In addition, 160 µL EDTA (0.5 M PH 8.0) and 80 µL NaCl (5 M) were added to dissolve pyrophosphate precipitates. Then, the RNA was precipitated by adding 900 µL isopropanol and incubating at −80°C for 2 h. The mixture was centrifuged at 21,000 xg at 4°C for 15 min, and the supernatant was discarded. The pellets were air-dried, resuspended in 150 µL RNAse-free water, 2X RNA loading dye was added to 1X (47.5% formamide, 0,01% bromophenol blue, 0.5 mM EDTA) and incubated at 70°C for 5 min. This mixture was then loaded onto a 12% denaturing urea polyacrylamide gel (8 M urea, 1X TBE (Sigma-Aldrich), 12% Bis-Tris acrylamide 19:1 (Carl Roth)) and run in 1X TBE at 300 V for 30 min. Afterward, the gel was covered in plastic wrap and placed on a fluor-coated cellulose TLC plate (Sigma-Aldrich) in a darkroom. The RNA bands were visualized using UV shadowing on the TLC plate at 254 nm.

The desired RNA band was cut out from the gel and shredded by passing the gel through two 3 mL syringes. 0.3 M NaOAc pH 5.2 (Invitrogen) was added to cover all gel pieces and incubated at −80°C overnight. Then, the small pieces were incubated at 37°C for 30 min and process of adding NaOAc and collecting the supernatant was repeated five times. The supernatants were filtered using a 0.22 µm syringe filter, precipitated with 70% ethanol, and incubated at −80°C overnight. On the next day, the mixture was centrifuged at 21,000 xg at 4°C for 30 min. The pellet was resuspended in RNAse-free water. Then, the RNA was purified using the Monarch RNA Cleanup Kit (500 µg, NEB). The concentration of the RNA was determined by measuring the absorbance at 260 nm using a NanoDrop, microvolume UV/Vis Spectrometer (Thermo Fisher). The RNA was stored at −80°C until further use.

### Capping of RNAs

The Vaccina capping cnzyme system (NEB) was used to generate the 5′ cap structure for the RNAs produced in the *in vitro* transcription reactions. For the cap(0) structure (m^7^GpppN-), up to 20 µg uncapped RNA was modified in a 40 µL reaction, containing 1 U/µL RiboLock (ThermoFisher), 1X capping buffer (NEB), 0.5 mM GTP (ThermoFischer), 0.2 mM S-adenosyl-methionine (SAM, NEB), and 2 µL of Vaccina capping enzyme (homemade, 3 mg/mL). For a cap(1) structure (m^7^GpppNm-) on the RNA, the reaction described above included another 2 µL of mRNA cap 2′-O-methyltransferase (50 U/µL, NEB). The capping reaction was incubated at 37°C for 4 h. Then, the RNA in the reaction was purified using the Monarch RNA Cleanup Kit (50 µg, NEB).

Capping was checked by loading 70 ng of the capped RNAs onto a 20% denaturing urea polyacrylamide gel. The gel was stained with SYBR Gold (1:10,000). The gels were scanned on the Typhoon FLA 9500 (GE Healthcare) for SYBR Gold.

### 3′-Cy5-labeling of RNAs

Up to 5 µg of RNA were used in a 20 µL reaction, containing additionally 0.5 mM ATP (Jena Bioscience), 50 µM Cy5-pCp (Jena Bioscience), 1X buffer (Jena Bioscience), 2 U/µL RiboLock (ThermoFisher), 1 µL T4 RNA ligase (Jena Bioscience). The mixture was incubated at 16°C overnight. The labeled RNA was purified using a Monarch RNA Cleanup Kit (10 µg, NEB).

### Endonuclease Activity Assay

For the endonuclease cleavage assay, 0.05 μM Cy5-labeled cap(1)-RNA (rGrArA rGrCrG rArGrA rArGrA rArCrA rCrArGrA rCrArG rCrArG rCrArG rArCrC rArGrG rC) was annealed to 0.05 μM of template DNA (GAT CAA GCT CAA GTA CTT AAG CCT GGT CTA TAC TAG TAC TGC C) in a thermocycler by heating to 72°C followed by cooling to 4°C at a rate of 0.1°C/s. 0.08 μM mammalian Pol II was added to the RNA: DNA hybrid and incubated at 30°C for 10 min. Then, 0.08 µM non-template DNA (GGC AGT ACT AGT ATT CTA GTA TTG AAA GTA CTT GAG CTT GAT C) was added and incubated at 30°C for 10 min.

Next, 0.12 μM of human elongation factors (DSIF) were added. Furthermore, 0.12 μM CAK and 1 mM ATP were added to generate phosphorylated Pol II. The mixture was incubated at 30°C for 30 min. After that, 0.04 μM viral FluPol with equimolar panhandle 5′ vRNA (/5Phos/rArGrU rArGrU rArArC rArArG rArG) and 3′ vRNA (rCrUrC rUrGrC rUrUrC rUrGrC rU) pre-incubated at 4°C were added. The reactions were incubated at 30°C, and samples were taken at 0, 10, and 60 min. These reactions occurred in 50 µL with a final buffer composition of 20 mM HEPES pH 7.4, 150 mM NaCl, 4% (v/v) glycerol, 3 mM MgCl_2_, 1 U/μL RiboLock (Thermo Fisher), and 1 mM TCEP.

The reactions were stopped by adding 1 µL of Proteinase K (NEB) to 7 µL of the sample and incubation at room temperature for 5 min. Then, 7 µL of 2X RNA Loading Dye (1X TBE, 3.6 M Urea, 0,01% bromophenol blue) was added to the sample. The samples were loaded onto 20% denaturing urea acrylamide gels and ran in 1X TBE buffer for 75 min at 300 V. The gels were scanned at the Typhoon FLA 9500 (GE Healthcare) for Cy5 fluorescence with PTM=750. This protocol was modified in the following way to check for Mg^2+^ dependence of the cap-snatching reaction during the sample. HEPES pH 7.4 was replaced by BICINE pH 8.5. The Mg^2+^ concentration was altered to 0.1 mM and 3 mM. The ATP concentration was changed to 0.01 mM and 1 mM to avoid complete chelating of Mg^2+^ by ATP. FluPol^E119D^ was used instead of wild type. The reaction was incubated at 30°C for 10 min, followed by 4°C overnight incubation, and then analyzed as described above.

### Quantification and Statistical Analysis of Endonuclease assays

The gels of the endonuclease activity assays were quantified using Fiji v2.9.0 ^60^. Therefore, the lanes were selected using rectangular selection masks. Then, the pixel intensities of each lane were plotted using the built-in gel-analysis functions. The intensity profile from each lane was examined, and individual bands could be distinguished as peaks. Vertical lines were drawn to delimit the peaks. The integrated intensities of each peak were measured and quantified as follows: the product band intensity was divided against the substrate band intensity. The procedure allows us to conclude a normalized cleavage ratio of the FluPol. The results were plotted using Graphpad Prism v9.4.1, indicating all individual data points as circles.

In Graphpad Prism, a Two-tailed paired parametric t-test with a 95% confidence interval was conducted between the indicated conditions. P-values are indicated in the figure.

### *In vitro* FluPol transcription activity assay

0.19 μM cap(1)-RNA (rGrArA rGrCrG rArGrA rArGrA rArCrA rCrArGrA rCrArG rCrArG rCrArG rArCrC rArGrG rC) was annealed to 0.19 μM of template DNA in a thermocycler by heating to 72°C followed by cooling to 4°C at a rate of 0.1°C/s. 0.31 μM mammalian Pol II was added to the RNA: DNA hybrid and incubated at 30°C for 10 min. Then, 0.31 μM non-template DNA was added and incubated at 30°C for 10 min. Next, 0.12 μM of DSIF were added. Furthermore, 0.50 μM CAK and 1 mM ATP were added to generate phosphorylated Pol II. The mixture was incubated at 30°C for 30 min. After that, 0.62 μM viral FluPol with modified panhandle vRNAs (3′vRNA with high G content, rCrUrG rUrGrU rGrCrC rUrCrU rGrCrU rUrCrU rGrCrU and 5’ vRNA /5Phos/rArGrU rArGrU rArArC rArArG rArG) pre-incubated at 4°C were added. Furthermore, 0.10 μM of CTP and GTP were added, as well as 0.77 µCi/µL α-^32^P-CTP. The reactions were incubated at 30°C for 2 h. Thse reactions occurred in 12.9 µL with a final buffer composition of 20 mM HEPES pH 7.4, 150 mM NaCl, 4% (v/v) glycerol, 3 mM MgCl_2_, 1 U/μL RiboLock (Thermo Fisher), and 1 mM TCEP.

The reactions were stopped by adding 1 µL of Proteinase K (NEB) to the sample and incubation at 37°C for 15 min. Then, 14 µL of 2X RNA Loading Dye (1X TBE, 3.6 M Urea, 0,01% bromophenol blue) was added to the sample. The samples were loaded onto 20% denaturing urea acrylamide gels and ran in 1X TBE buffer for 75 min at 300 V. The gels were incubated for 2 h on a phosphorus screen. The screen was scanned at the Typhoon FLA 9500 (GE Healthcare) with PTM=800.

### Analytical Gel Filtration on Äkta µ

For an assembly in a 50 µL reaction, 42.75 pmol RNA was annealed to 42.75 pmol template DNA as described for the endonuclease assay. 28.5 pmol mammalian Pol II was added to the RNA: DNA scaffold, followed by 57 pmol of non-template DNA, and incubated at 30°C for 10 min after each addition. Next, 0.8 μM CAK, 1 mM ATP, and 57 pmol human transcription elongation factors were added and incubated at 30°C for 30 min. The CAK was omitted for the non-phosphorylation assays. Then, pre-mixed 57 pmol viral FluPol (endonuclease inactive version PA^E119D^) with equimolar panhandle 5′ vRNA (/5Phos/rArGrU rArGrU rArArC rArArG rArG) and 3′ vRNA (rCrUrC rUrGrC rUrUrC rUrGrC rU) were added to the mix. Lastly, the reaction was incubated at 30°C for an additional 10 min. The final buffer composition was 50 mM Bicine pH 8.5 at 4°C, 150 mM NaCl, 4% (v/v) glycerol, 3 mM MgCl_2_, and 1 mM TCEP.

The fully formed complex was centrifuged at 21,000 xg at 4°C for 10 min. The supernatant was injected onto a Superose 6 Increase 3.2/300 column (Cytiva) and ran in SEC buffer (20 mM Bicine pH 8.5 at 4°C, 150 mM NaCl, 4% (v/v) glycerol, 3 mM MgCl_2_, 1 mM TCEP) on an ÄKTAmicro (GE Healthcare) system. The absorbances at 280 nm (protein) and 260 nm (RNA/DNA) were measured. The absorbance data were plotted using GraphPad Prism v9.4.1. The main elution fractions were analyzed by SDS-PAGE.

### Western Blot

Samples of the peak fractions were collected to compare the presence of FluPol in the Pol II containing fractions, mixed with 4X SDS-loading dye (ThermoFisher), and stored at −20 °C until analysis.

The samples were run on one SDS-PAGE (NuPAGE 4-12% Bis-Tris, Invitrogen) in 1X MES buffer (Invitrogen). The gel was then blotted onto a nitrocellulose membrane (GE Healthcare) using a wet-blot system (ThermoFisher) in NuPAGE transfer buffer (Invitrogen). The blot was then blocked for 1 h at room temperature with 5% (w/v) milk powder in PBS-T. Then, the membrane was cut horizontally at the 50 kDa line. The upper half was incubated overnight with a rabbit anti-Strep antibody (1:1,000 dilution; ab76949, Abcam) against the StrepTag II on the FluPol. The lower half was incubated with a rabbit anti-RPB3 polyclonal (1:2000 dilution; A303-771A, Bethyl) as a loading control.

The following day, the membranes were washed 3x 1 min and 3x 10 min with PBS-T and incubated with an anti-rabbit antibody coupled to HRP (1:1000; homemade) in PBS-T with 5% milk powder. Then, the membrane was washed three times with PBS-T for 10 min, developed with SuperSignal West Pico Substrate (Thermo Fisher), and scanned using a ChemoCam Advanced Fluorescence imaging system (Intas Science Imaging).

To assess steady-state levels of A/WSN/33-derived PA and PB2 proteins, total lysates of HEK-293T cells transfected with the corresponding pcDNA3.1 expression plasmid were prepared in Laemmli buffer. Proteins were separated by SDS-PAGE using NuPAGE™ 4-12% Bis-Tris gels (Invitrogen) and transferred to nitrocellulose membranes which were incubated with primary antibodies directed against PA (GTX125932 - 1:5,000), PB2 (GTX125925 - 1:5,000), or Tubulin (Sigma-Aldrich T5168 - 1:10,000) and subsequently with HRP-tagged secondary antibodies (Sigma Aldrich, A9044 and A9169, 1:10,000). Membranes were developed with the ECL2 substrate according to the manufacturer′s instructions (Pierce) and chemiluminescence signals were acquired using the ChemiDoc imaging system (Bio-Rad). Uncropped gels are provided as a source data file.

### Sample Preparation for Cryo-EM

First, 180 pmol cap(1)-RNA was annealed to 180 pmol 5′-Cy5-labeled template DNA, as stated previously. 120 pmol mammalian Pol II was added to the RNA-DNA scaffold and incubated at 30°C for 10 min. Then, 240 pmol of non-template was added and kept at 30°C for 10 min. Next, 1 μM CAK, 1 mM ATP, and 240 pmol human transcription elongation factors were added and incubated at 30°C for 30 min. Lastly, pre-mixed 240 pmol viral FluPol (endonuclease inactive version PA^E119D^) with equimolar 5′/3′-vRNAs was added to the mix and incubated at 30°C for 10 min. The 3′-vRNA was ATTO532-labeled on the 5′-end. The complex was assembled in a buffer containing 50 mM Bicine pH 8.5 at 4°C, 150 mM NaCl, 4% (v/v) glycerol, 0.1 mM MgCl_2_ (3 mM MgCl_2_ for post-cleavage conformation, 0.1 mM MgCl_2_ for pre-cleavage conformation), and 1 mM TCEP in a volume of 150 µL. The fully formed complex was centrifuged at 21,000 xg at 4°C for 10 min.

The sample was loaded on a continuous 10-40% glycerol gradient containing assembly buffer components. The heavy solution contained additionally 0.1% (v/v) glutaraldehyde.

The gradient was centrifuged at 33,000 rpm in a SW60 rotor (Beckman Coulter) at 4°C for quenched by adding 100 mM Tris-HCl pH 8.0 at 4°C. Fractions were analyzed by NativePAGE 3-12% (Bis-Tris, Invitrogen) run at 4°C. The gel was then scanned for Cy5 and ATTO532 signals, followed by Coomassie staining.

Then, the complex containing fractions were dialyzed against 20 mM Tris pH 8 at 20°C, 20 mM Bicine pH 8.5 at 4°C, 100 mM NaCl, 4% (v/v) glycerol, 0.1 mM MgCl_2_ (3 mM MgCl_2_ for post-cleavage conformation, 0.1 mM MgCl_2_ for pre-cleavage conformation), and 1 mM TCEP using a 20 kDa Slide-A-Lyzer™ MINI device (Thermo Fisher) at 4°C for 4 h. Onto the sample was a continuous carbon film of roughly 3 nm floated for 5 min. The carbon was then fished with a glow-discharged holey carbon grid (Quantifoil R3.5/1, copper, mesh 200). 4 µL of dialysis buffer was added to the grid, and the grid was placed in a Vitrobot Mark IV (Thermo Fisher) under 100% humidity at 4°C. The grids were then blotted using Whatman paper with a blot force of 5 for 5 s and directly plunge-frozen in liquid ethane.

### Cryo-EM analysis and image processing

A Titan Krios G2 transmission electron microscope (FEI) operated at 300 keV, equipped with a GIF BioQuantum energy filter (Gatan) and a K3 summit direct detector was used to acquire cryo-EM data. Data acquisition was performed at a pixel size of 1.05 Å/pixel using Serial EM, corresponding to a nominal magnification of 81,000X in nanoprobe EFTEM mode.

The pre-cleavage dataset was collected in 5 batches. A total of 60,032 movie stacks were collected. Each movie contained 40 frames and was acquired in counting mode over 1.95 s. The defocus was set to values between −0.1 to −2.0 µm. The dose rate was 20.48 e^−^ per Å^2^ per s, leading to a total dose of 39.94 e^−^ per Å^2^.

The post-cleavage dataset was collected in 3 batches. A total of 20,509 movie stacks were collected. Each movie contained 40 frames and was acquired in counting mode over 2.4 s. The defocus was set to values between −0.1 to −2.0 µm. The dose rate was 18.34 e^−^ per Å^2^ per s, leading to a total dose of 40 e^−^ per Å^2^.

Data preprocessing, including stacking, contrast transfer function (CTF) estimation, and dose-weighting, was done using Warp^61^. In Warp, particles were also picked using an on this data set trained version of the neural network BoxNet2.

For the post-cleavage dataset, 11,935,228 particles were extracted in five batches in RELION-3.1.0^62^ using a binning factor of four. The box size of the particles was set to 112 pixels with a pixel size of 4.2 Å/px. The particles were then imported into cryoSPARC v4.3.1^63^. In cryoSPARC, particles that do not align were removed, as well as particles that do not contain Pol II using 3D heterogeneous refinements. The 1,975,313 particles that contain Pol II were transferred to RELION and extracted with a box size of 448 px and a pixel size of 1.05 Å/px. These particles were refined using a mask around the Pol II core, followed by Bayesian polishing and CTF refinement for beam tilt and per-particle defocus values. The particles were reloaded into cryoSPARC, combined into 3 datasets, followed by 1 round of heterogeneous refining for FluPol occupancy. Then, the datasets were individually non-uniformly refined, locally refined onto the FluPol, and 3D classified. The data sets were merged, locally refined for FluPol, and 2 times 3D classified. From a final dataset of 63,230 particles, focus refinements on FluPol, Pol II core, Pol II stalk, and the interface were performed.

For the pre-cleavage dataset, 6,423,874 particles were extracted in three batches in RELION-3.1^62^ using a pixel size of 4.2 Å/px and a box size of 112 pixels. The particles were then imported into cryoSPARC v4.3.1^63^. In cryoSPARC, particles that do not align were removed, as well as particles that do not contain Pol II using 3D heterogeneous refinements. The 1,937,625 particles that contain Pol II were transferred to RELION and extracted with a box size of 448 px and a pixel size of 1.05 Å/px. These particles were refined using a mask around the Pol II core, followed by Bayesian polishing and CTF refinement for beam tilt and per-particle defocus values. The particles were focused refined, and classified on the Pol II core, taking only the particles of the class with good-looking Pol II. These particles were globally classified for FluPol occupancy and then focussed classified on FluPol for well-aligning FluPol particles. This final particle set of 369,858 particles was focused refined on Pol II, CTF refined, and Bayesian polished. Focus refinements for Pol II core and FluPol were performed based on the obtained consensus refinement.

### Model building

For both structures, initial models of *Sus scrofa domesticus* Pol II (PDB: 7B0Y^64^), SPT5 KOW2, KOW3, KOWx-4 and KOW5 domains (PDB: 5OIK and 5OHO^34^) and FluPolA/H7N9 (PDB:7QTL ^36^)) were rigid body fitted in ChimeraX 1.6.1^65^ using the consensus refinement. The RNA and DNA were manually adjusted in Coot^66^ to fit the sequences used in this study. As the density of the RNA in the endonuclease site is not well enough resolved to call a sequence, we modeled the sequence according to the biochemistry. The linker between KOWx-4 and KOW5 was manually built as well, assuming that the best visible amino acid at the G1 nucleotide is the first phenylalanine of the linker. This model, the focused maps, and the consensus map were loaded into ISOLDE 1.6.0^67^. The focused maps were aligned to the consensus map in ChimeraX. In ISOLDE, Molecular Dynamic simulation was performed using the starting model restrains. Then, the individual protein components were subjected to Real Space Refinement in PHENIX^68^ and manual curation in Coot.

For the pre-cleavage conformation, Pol II and KOW5 were refined against the focused map for Pol II. FluPol was refined against the focused map for the FluPol. KOWx-4 was refined against the consensus map.

Pol II (except RPB4 and RPB7) was refined against the Pol II-focused map for the post-cleavage state. SPT5 KOW2, KOW3, KOWx-4, RPB4, and RPB7 were refined against the stalk-focused map. FluPol was refined against the focused map for the FluPol. Then, the Pol II elongation complex components and the FluPol were rigid body docked into the consensus map in ChimeraX before manually checking interface residues in Coot using the consensus map. The density for SPT4 and SPT4 NGN and KOW1 domain is not well resolved, so the consensus map was lowpass filtered to 6 Å. A deposited model (PDB: 5OIK for pre-cleavage and 7YCX^69^ for post-cleavage) for these domains was rigid body fitted into this filtered map using ChimeraX. Interfaces were checked for major clashes. Clashing residues without density were modified in Coot using the most likely, non-clashing rotamer.

To determine the range of possible RNA lengths, a series of FluPol structures was modeled in Coot by cropping nucleotides in the less-resolved space between cap-binding domain and endonuclease domain. Then, these structures were loaded into ISOLDE and the RNA was real-space refined. The lower limit was defined as when ISOLDE shifted the RNA through the endonuclease domain. The upper limit was determined by incrementally increasing the RNA length in Coot, refined in ISOLDE, and visually inspected until the obtained model deviated from an expected linear RNA geometry.

### Cell-based minigenome assay

The plasmids and procedure used for minigenome assays are described in ^42^. The primers used for mutagenesis of the PB2 and PA plasmids can be provided upon request. Briefly, HEK-293T cells were co-transfected with plasmids encoding the vRNP protein components (PB2, PB1, PA, NP), a pPolI-Firefly plasmid encoding a negative-sense viral-like RNA expressing the Firefly luciferase and the pTK-Renilla plasmid (Promega) as an internal control. Mean relative light units (RLUs) produced by the Firefly and Renilla luciferase, reflecting the viral polymerase activity and transfection efficiency, respectively, were measured using the Dual-Glo Luciferase Assay System (Promega) on a Centro XS LB960 microplate luminometer (Berthold Technologies, MikroWin Version 4.41) at 24 hours post-transfection (hpt). Firefly luciferase signals were normalised with respect to Renilla luciferase signals. Three independent experiments (each in technical duplicates) were performed.

### Selection of interface residues for mutational analysis

First, a list of 38 amino residues at the interfaces was generated, see **Table 2**. In SnapGene, 7.0.1 two MUSCLE alignments were performed for PA and PB2 (**ED Figure 4a-b**). Each alignment contained sequences of 6 influenza A viruses, 2 influenza B viruses, and one influenza C and D virus. Sequences of the following strains were used: A/Zhejiang/HZ1/2013 (H7N9), A/WSN/1933 (H1N1), A/California/04/2009 (H1N1), A/California/04/2009 (H1N1), A/Victoria/3/1975 (H3N2), A/Little-yellow-shouldered-bat/Guatemala/2010 (H17N10), B/Lee/1940, B/Memphis/13/2003, C/Johannesburg/1/1966, D/Bovine/Minnesota/628/2013. The 16 residues with a MUSCLE score of above 50 were considered conserved. These amino acids were mutated to alanine. All mutants were checked for the expression level. The mutant Y131A showed a reduced expression level and was consequently excluded for further analysis. We tested furthermore the following double and triple mutants: PA S140A, E141A; PB2 R375A, R380A; PB2 D466A, T468A, S470A. From these mutants, only PB2 R375A, R380A had wildtype expression levels.

To investigate the evolutionary conservation of the binding interfaces between mammals and birds, four mammalian species, four bird species and *C. elegans* were used. Sequences were identified using the BLAST algorithm of Uniprot using the selected species as a search target. To select for the bird species, the human RPB1 sequence was blasted against all avian protein sequences available in Uniprot. Only four bird species had full-length annotated RPB1. These species were used as a search filter while blasting human RPB3, RPB11 and SPT5. A list of all Uniprot sequence IDs is available upon request. The obtained sequences were aligned in Snapgene using the ClustalOmega algorithm. Alignments are depicted in **ED Figure 4d-g**.

## References

1. Plotch, S. J., Bouloy, M., Ulmanen, I. & Krug, R. M. A unique cap(m7GpppXm)-dependent influenza virion endonuclease cleaves capped RNAs to generate the primers that initiate viral RNA transcription. Cell 23, 847–858 (1981).

2. Lukarska, M. et al. Structural basis of an essential interaction between influenza polymerase and Pol II CTD. Nature 541, 117–121 (2017).

3. Mahy, B. W. J., Hastie, N. D. & Armstrong, S. J. Inhibition of Influenza Virus Replication by α-Amanitin: Mode of Action. Proc. Natl. Acad. Sci. 69, 1421–1424 (1972).

4. te Velthuis, A. J. W. & Fodor, E. Influenza virus RNA polymerase: insights into the mechanisms of viral RNA synthesis. Nat. Rev. Microbiol. 14, 479–493 (2016).

5. Krischuns, T., Lukarska, M., Naffakh, N. & Cusack, S. Influenza Virus RNA-Dependent RNA Polymerase and the Host Transcriptional Apparatus. Annu. Rev. Biochem. (2021) doi:10.1146/annurev-biochem-072820-100645.

6. Iuliano, A. D. et al. Estimates of global seasonal influenza-associated respiratory mortality: a modelling study. The Lancet 391, 1285–1300 (2018).

7. WHO. Influenza (seasonal) fact sheet 2023. https://www.who.int/en/news-room/fact-sheets/detail/influenza-(seasonal) (2023).

8. Taubenberger, J. K. & Morens, D. M. 1918 Influenza: the Mother of All Pandemics. Emerg. Infect. Dis. 12, 15–22 (2006).

9. Krammer, F. et al. Influenza. Nat. Rev. Dis. Primer 4, 3 (2018).

10. Eisfeld, A. J. et al. Pathogenicity and transmissibility of bovine H5N1 influenza virus. Nature (2024) doi:10.1038/s41586-024-07766-6.

11. Mallapaty, S. Bird flu could become a human pandemic. How are countries preparing? Nature d41586–024-02237–4 (2024) doi:10.1038/d41586-024-02237-4.

12. CDC. Current H5N1 Bird Flu Situation in Dairy Cows. https://www.cdc.gov/bird-flu/situation-summary/mammals.html#cdc_generic_section_5-resouces.

13. Chou, Y. et al. One influenza virus particle packages eight unique viral RNAs as shown by FISH analysis. Proc. Natl. Acad. Sci. 109, 9101–9106 (2012).

14. Herz, C., Stavnezer, E., Krug, R. M. & Gurney, T. Influenza virus, an RNA virus, synthesizes its messenger RNA in the nucleus of infected cells. Cell 26, 391–400 (1981).

15. Wandzik, J. M. et al. A Structure-Based Model for the Complete Transcription Cycle of Influenza Polymerase. Cell 181, 877–893.e21 (2020).

16. Pflug, A., Guilligay, D., Reich, S. & Cusack, S. Structure of influenza A polymerase bound to the viral RNA promoter. Nature 516, 355–360 (2014).

17. Ramanathan, A., Robb, G. B. & Chan, S.-H. mRNA capping: biological functions and applications. Nucleic Acids Res. 44, 7511–7526 (2016).

18. Plotch, S. J., Tomasz, J. & Krug, R. M. Absence of Detectable Capping and Methylating Enzymes in Influenza Virions. J. Virol. 28, 75–83 (1978).

19. Kouba, T., Drncová, P. & Cusack, S. Structural snapshots of actively transcribing influenza polymerase. Nat. Struct. Mol. Biol. 26, 460–470 (2019).

20. Sikora, D., Rocheleau, L., Brown, E. G. & Pelchat, M. Deep sequencing reveals the eight facets of the influenza A/HongKong/1/1968 (H3N2) virus cap-snatching process. Sci. Rep. 4, 6181 (2014).

21. Osman, S. & Cramer, P. Structural Biology of RNA Polymerase II Transcription: 20 Years On. Annu. Rev. Cell Dev. Biol. 36, 1–34 (2020).

22. Akhtar, Md. S., et al. TFIIH Kinase Places Bivalent Marks on the Carboxy-Terminal Domain of RNA Polymerase II. Mol. Cell 34, 387–393 (2009).

23. Velychko, T. et al. CDK7 kinase activity promotes RNA polymerase II promoter escape by facilitating initiation factor release. Mol. Cell S1097–2765(24)00400–3 (2024)

24. Zhan, Y., Grabbe, F., Oberbeckmann, E., Dienemann, C. & Cramer, P. Three-step mechanism of promoter escape by RNA polymerase II. Mol. Cell 84, 1699–1710.e6 (2024).

25. Core, L. & Adelman, K. Promoter-proximal pausing of RNA polymerase II: a nexus of gene regulation. Genes Dev. 33, 960–982 (2019).

26. Garg, G. et al. Structural insights into human co-transcriptional capping. Mol. Cell 83, 2464–2477.e5 (2023).

27. Tsukamoto, Y. et al. Inhibition of cellular RNA methyltransferase abrogates influenza virus capping and replication. Science 379, 586–591 (2023).

28. Liu, Y. et al. The Crystal Structure of the PB2 Cap-binding Domain of Influenza B Virus Reveals a Novel Cap Recognition Mechanism. J. Biol. Chem. 290, 9141–9149 (2015).

29. Chan, A. Y., Vreede, F. T., Smith, M., Engelhardt, O. G. & Fodor, E. Influenza virus inhibits RNA polymerase II elongation. Virology 351, 210–217 (2006).

30. Engelhardt, O. G., Smith, M. & Fodor, E. Association of the Influenza A Virus RNA-Dependent RNA Polymerase with Cellular RNA Polymerase II. J. Virol. 79, 5812–5818 (2005).

31. Lukarska, M. & Cusack, S. Structural and functional characterization of the interaction between influenza polymerase and the cellular transcription machinery, Caractérisation structurale et fonctionnelle de l’intéraction entre la polymérase d’influenza et la machinerie cellulaire de transcription. (Université Grenoble Alpes, 2018) https://theses.hal.science/tel-02954348v1/file/LUKARSKA_2018_archivage.pdf.

32. Krischuns, T. et al. Type B and type A influenza polymerases have evolved distinct binding interfaces to recruit the RNA polymerase II CTD. PLOS Pathog. 18, e1010328 (2022).

33. Bradel-Tretheway, B. G. et al. Comprehensive Proteomic Analysis of Influenza Virus Polymerase Complex Reveals a Novel Association with Mitochondrial Proteins and RNA Polymerase Accessory Factors. J. Virol. 85, 8569 (2011).

34. Bernecky, C., Plitzko, J. M. & Cramer, P. Structure of a transcribing RNA polymerase II–DSIF complex reveals a multidentate DNA–RNA clamp. Nat. Struct. Mol. Biol. 24, 809– 815 (2017).

35. Chen, Y. et al. Human infections with the emerging avian influenza A H7N9 virus from wet market poultry: clinical analysis and characterisation of viral genome. The Lancet 381, 1916–1925 (2013).

36. Kouba, T. et al. Direct observation of backtracking by influenza A and B polymerases upon consecutive incorporation of the nucleoside analog T1106. Cell Rep. 42, 111901 (2023).

37. Song, M.-S. et al. Identification and characterization of influenza variants resistant to a viral endonuclease inhibitor. Proc. Natl. Acad. Sci. U. S. A. 113, 3669–3674 (2016).

38. Kumar, G., Cuypers, M., Webby, R. R., Webb, T. R. & White, S. W. Structural insights into the substrate specificity of the endonuclease activity of the influenza virus cap-snatching mechanism. Nucleic Acids Res. 49, 1609–1618 (2021).

39. Bernecky, C., Herzog, F., Baumeister, W., Plitzko, J. M. & Cramer, P. Structure of transcribing mammalian RNA polymerase II. Nature 529, 551–554 (2016).

40. Koppstein, D., Ashour, J. & Bartel, D. P. Sequencing the cap-snatching repertoire of H1N1 influenza provides insight into the mechanism of viral transcription initiation. Nucleic Acids Res. 43, 5052–5064 (2015).

41. Stark, H. GraFix: Stabilization of Fragile Macromolecular Complexes for Single Particle Cryo-EM. in Methods in Enzymology vol. 481 109–126 (Elsevier, 2010).

42. Krischuns, T. et al. The host RNA polymerase II C-terminal domain is the anchor for

43. Keown, J. et al. Structural and functional characterization of the interaction between the influenza A virus RNA polymerase and the CTD of host RNA polymerase II. J. Virol. 98, e00138–24 (2024).

44. Fan, H. et al. Structures of influenza A virus RNA polymerase offer insight into viral genome replication. Nature 573, 287–290 (2019).

45. Fong, N., Sheridan, R. M., Ramachandran, S. & Bentley, D. L. The pausing zone and control of RNA polymerase II elongation by Spt5: Implications for the pause-release model. Mol. Cell 82, 3632–3645.e4 (2022).

46. Reich, S., Guilligay, D. & Cusack, S. An *in vitro* fluorescence based study of initiation of RNA synthesis by influenza B polymerase. Nucleic Acids Res. gkx043 (2017) doi:10.1093/nar/gkx043.

47. Reich, S. et al. Structural insight into cap-snatching and RNA synthesis by influenza polymerase. Nature 516, 361–366 (2014).

48. Vos, S. M., Farnung, L., Urlaub, H. & Cramer, P. Structure of paused transcription complex Pol II–DSIF–NELF. Nature 560, 601–606 (2018).

49. Vos, S. M. et al. Structure of activated transcription complex Pol II–DSIF–PAF–SPT6. Nature 560, 607–612 (2018).

50. Fianu, I. et al. Structural basis of Integrator-dependent RNA polymerase II termination. Nature 629, 219–227 (2024).

51. Zimmer, J. T., Rosa-Mercado, N. A., Canzio, D., Steitz, J. A. & Simon, M. D. STL-seq reveals pause-release and termination kinetics for promoter-proximal paused RNA polymerase II transcripts. Mol. Cell 81, 4398–4412.e7 (2021).

52. Gressel, S. et al. CDK9-dependent RNA polymerase II pausing controls transcription initiation. eLife 6, e29736 (2017).

53. Bier, K., York, A. & Fodor, E. Cellular cap-binding proteins associate with influenza virus mRNAs. Journal of General Virology vol. 92 1627–1634 (2011).

54. Lu, H. et al. Recent advances in the development of protein–protein interactions modulators: mechanisms and clinical trials. Signal Transduct. Target. Ther. 5, 213 (2020).

55. Abramson, J. et al. Accurate structure prediction of biomolecular interactions with AlphaFold 3. Nature 630, 493–500 (2024).

56. Krishna, R. et al. Generalized biomolecular modeling and design with RoseTTAFold All-Atom. Science 384, eadl2528 (2024).

57. Zeng, Y., Zhang, H.-W., Wu, X.-X. & Zhang, Y. Structural basis of exoribonuclease-mediated mRNA transcription termination. Nature 628, 887–893 (2024).

58. Boehning, M. et al. RNA polymerase II clustering through carboxy-terminal domain phase separation. Nat. Struct. Mol. Biol. 25, 833–840 (2018).

59. Lu, C. & Li, P. Preparation of Short RNA by In Vitro Transcription. in Recombinant and In Vitro RNA Synthesis (ed. Conn, G. L.) vol. 941 59–68 (Humana Press, Totowa, NJ, 2013).

60. Schindelin, J., et al. Fiji: an open-source platform for biological-image analysis. Nat. Methods 9, 676–682 (2012).

61. Tegunov, D. & Cramer, P. Real-time cryo-electron microscopy data preprocessing with Warp. Nat. Methods 16, 1146–1152 (2019).

62. Zivanov, J. et al. New tools for automated high-resolution cryo-EM structure determination in RELION-3. eLife 7, e42166 (2018).

63. Punjani, A., Rubinstein, J. L., Fleet, D. J. & Brubaker, M. A. cryoSPARC: algorithms for rapid unsupervised cryo-EM structure determination. Nat. Methods 14, 290–296 (2017).

64. Zhang, S. et al. Structure of a transcribing RNA polymerase II-U1 snRNP complex. Science 371, 305–309 (2021).

65. Pettersen, E. F. et al. UCSF CHIMERAX: Structure visualization for researchers, educators, and developers. Protein Sci. 30, 70–82 (2021).

66. Emsley, P., Lohkamp, B., Scott, W. G. & Cowtan, K. Features and development of Coot. Acta Crystallogr. D Biol. Crystallogr. 66, 486–501 (2010).

67. Croll, T. I. *ISOLDE*: a physically realistic environment for model building into low-resolution electron-density maps. Acta Crystallogr. Sect. Struct. Biol. 74, 519–530 (2018).

68. Liebschner, D. et al. Macromolecular structure determination using X-rays, neutrons and electrons: recent developments in *Phenix*. Acta Crystallogr. Sect. Struct. Biol. 75, 861– 877 (2019).

69. Zheng, H. et al. Structural basis of INTAC-regulated transcription. Protein Cell 14, 698–702 (2023).

